# PEPSDI: Scalable and flexible inference framework for stochastic dynamic single-cell models

**DOI:** 10.1101/2021.07.01.450748

**Authors:** Sebastian Persson, Niek Welkenhuysen, Sviatlana Shashkova, Samuel Wiqvist, Patrick Reith, Gregor W. Schmidt, Umberto Picchini, Marija Cvijovic

## Abstract

Mathematical modelling is an invaluable tool to describe dynamic cellular processes and to rationalise cell-to-cell variability within the population. This requires statistical methods to infer unknown model parameters from dynamic, multi-individual data accounting for heterogeneity caused by both intrinsic and extrinsic noise. Here we present PEPSDI, a scalable and flexible framework for Bayesian inference in state-space mixed-effects stochastic dynamic single-cell models. Unlike previous frameworks, PEPSDI imposes a few modelling assumptions when inferring unknown model parameters from time-lapse data. Specifically, it can infer model parameters when intrinsic noise is modelled by either exact or approximate stochastic simulators, and when extrinsic noise is modelled by either time-varying, or time-constant parameters that vary between cells. This allowed us to identify hexokinase activity as a source of extrinsic noise, and to deduce that sugar availability dictates cell-to-cell variability in the budding yeast *Saccharomyces cerevisiae* SNF1 nutrient sensing pathway.

---

Traditionally, investigations in the life sciences have focused on a population “ensemble average” level. On one side, such population approach reduces noise from atypical cells. However, any cellular population is in general heterogeneous, with a range of different physical, chemical, and biological properties. Thus, population methods smooth out and hence miss biologically relevant cell-to-cell variability [1]. For example, such approaches will overlook drug-resistant bacteria or cancer cells in a general cell populations. Furthermore, cell-to-cell variability plays an important role in the decision making of a population, such as quick adaptation to fluctuating environments [2]. The only way to identify all biologically relevant processes, and thus describe cell heterogeneity, is to study the whole population cell-by-cell [3].

To study life processes occurring in individual cells within the population, fluorescent time-lapse microscopy can be employed to track proteins in multiple cells over time [1]. Ideally, this can give a view of cell heterogeneity, and potentially help elucidate cellular reaction dynamics. However, to learn more from acquired data, dynamic modelling naturally complements time-lapse microscopy, and aids in deducing sources of cell-to-cell variability [4, 5, 6]. But to fully exploit dynamic modelling, unknown model parameters must typically be inferred/estimated from data [7]. This is non-trivial to perform from single-cell time-lapse data, mainly because models describing individual cells must account for cell-to-cell variability caused by both intrinsic (e.g. variations in chemical reactions) and extrinsic (e.g. variability in protein concentrations) noise [8].

Several inference methods exist that, to various degrees, account for cell-to-cell variability when deducing model parameters from time-lapse data. In common, they allow extrinsic noise to be modelled by letting model parameters, e.g protein synthesis rates, vary between cells [4, 6, 9, 10]. Methods based on ordinary differential equations (ODEs) [6] further assume that intrinsic noise is negligible. On the other side, the dynamic prior propagation (DPP) [4] and the stochastic differential equation mixed-effects models (SDEMEM) [9, 10] encode intrinsic noise via exact [11], or approximate [12] stochastic simulators, respectively.

Although useful, current inference methods have drawbacks. The fact that ODE based methods assume intrinsic noise to be negligible is often hard to justify. The DPP method imposes multiple model assumptions, such as time invariant rate constants. The SDEMEM methods employ approximate simulators to model intrinsic noise, that can be inaccurate when few molecules control the dynamics [13]. Overall, available frameworks only address specific questions. Moreover, there are scenarios where all existing methods are inadequate. For example, when studying a gene expression model with low numbers of molecules and a time-varying transcription rate using available inference approaches will impose unrealistic model assumptions, potentially leading to incorrect model prediction [14].

To fully exploit the power and facilitate usage of stochastic single-cell dynamic models, we propose, for the first time, a flexible Bayesian inference framework for stochastic dynamic mixed-effects models. By building upon a Bayesian inference framework originally thought for SDEMEMs [10], we introduce a novel, computationally more efficient inference method, that is capable of inferring unknown model parameters when intrinsic noise is modelled by either exact [11, 15], or approximate [12, 13] stochastic simulators. Moreover, by leveraging on the state-of-the-art statistical methods [16, 17], our framework allows for large flexibility in how extrinsic noise is modelled. Using synthetic examples, we show how this flexibility facilitates understanding of a stochastic gene expression model regulated by an extrinsic time-varying signal and cellular pathways where intrinsic noise causes cells to migrate between states. Further, by combining time-lapse microscopy with microfluidics, we employ our inference framework to distinguish between multiple network structures and identify sources of cell heterogeneity in the budding yeast *Saccharomyces cerevisiae* SNF1 nutrient sensing pathway.

## Results

### Inference framework for stochastic dynamic single-cell models

We developed a flexible modelling framework, PEPSDI (**P**articles **E**ngine for **P**opulation **S**tochastic **D**ynam**I**cs), that infers unknown model parameters from dynamic data for single-cell dynamic models that account for both intrinsic and extrinsic noise (Fig. 1). The latter can be modelled by letting parameters believed to vary between cells, such as protein translation rates, follow a probability distribution. Furthermore, the model parameters can incorporate extrinsic time variant signals, such as the circadian clock [18], and measured extrinsic data, such as cell volume. Intrinsic noise can be modelled by multiple stochastic simulators, specifically the exact SSA (Gillespie) and Extrande simulators [11, 15], and the approximate tau-leaping [13] and Langevin simulators [12]. Hence, PEPSDI is applicable for gene expressions models with low numbers of molecules [19], and signalling models where large numbers of molecules can make exact simulators unfeasible [13]. This framework can further infer the strength of the measurement error, and is suitable when either all, or a subset of the model components are observed. More formally, PEPSDI produces Bayesian inference for state-space models with latent dynamics incorporating mixed-effects, that is state-space mixed-effects model (SSMEM). It builds upon the schemes previously proposed for SDEMEMs [10].

**Figure 1:**
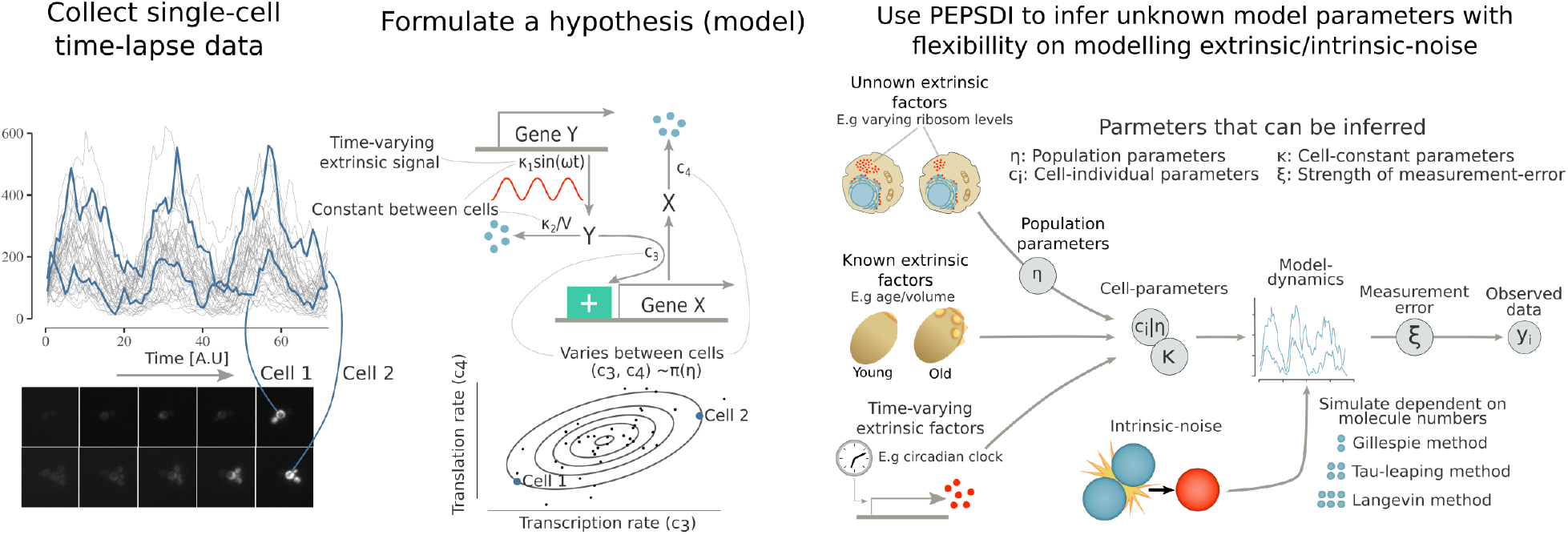
PEPSDI: A Bayesian inference framework for single-cell stochastic dynamic models. Single-cell time-lapse data obtained via fluorescent microscopy often exhibits considerable cell-to-cell variability (left). Dynamic modelling (middle) can help elucidate both the reaction dynamics, and sources of cell-to-cell variability behind such data. PEPSDI (right) is a flexible inference framework for dynamic stochastic single-cell models that imposes few model assumptions. For example, extrinsic noise can be modelled by letting cell-individual quantities i) be modelled probabilistically as **c**^(*i*)^ ~ *π*(**c**^(*i*)^|***η***) (unknown extrinsic factors), ii) be combined with known extrinsic data (known extrinsic factors), and iii) be time-variant (time varying extrinsic factors). Furthermore, PEPSDI includes multiple stochastic algorithms for modelling intrinsic noise, and assumes that the observed data **y**^(*i*)^ is acquired with a measurement error. Overall, based on the observed data PEPSDI can infer the cell individual parameters **c**^(*i*)^, the cell constant parameters ***κ***, the population parameters ***η*** and the strength of the measurement error ***ξ***.

From the methodological point of view, PEPSDI is a Gibbs sampler targeting the full posterior distribution of all unknowns. To allow large flexibility in the model construction, some of the Gibbs-steps can be targeted using Hamiltonian Monte Carlo (HMC) [17]. For example, it can be assumed that the synthesis and breakdown rates of a protein follow a log-normal distribution and if correlation between the rates is suspected, *a-priori*, the HMC sampler permits efficient inference of the correlation [20]. For the Gibbs-steps where the likelihood function is intractable, PEPSDI uses a pseudo-marginal approach employing particle filters [16]. This enables the user to select from a wide range of stochastic simulators [13]. Furthermore, for computational efficiency we employ, when possible, correlated particle filters [21], and tune the parameters proposal distributions using adaptive algorithms [22, 23, 24].

PEPSDI can be run with two options. The default is to allow cell constant parameters to vary weakly between cells, and the second option is to keep cell constant parameter fixed. The former speeds up the inference by a factor of 35 compared to the latter (Supplementary Fig. S3), and larger speed ups are likely attainable when the number of observed cells increase.

PEPSDI is written in Julia [25], and is available on GitHub (https://github.com/cvijoviclab/PEPSDI). To encourage usage of our framework, the provided code is flexible with regards to the model structure and all examples are available as notebooks (https://github.com/cvijoviclab/PEPSDI/tree/main/Code/Examples). Guidelines on how to run the inference schemes are presented in Supplementary Note 3, and a tutorial in Supplementary Note 4.

### Application to simulated circadian clock gene expression model

We applied the developed inference framework on simulated data from a simple circadian clock [18] gene expression model (Fig. 2a). The circadian clock was modelled as a sine function regulating the transcription activity, causing the protein levels to oscillate (Fig. 2b). Moreover, to simulate strong intrinsic noise, the numbers of molecules were kept low.

**Figure 2:**
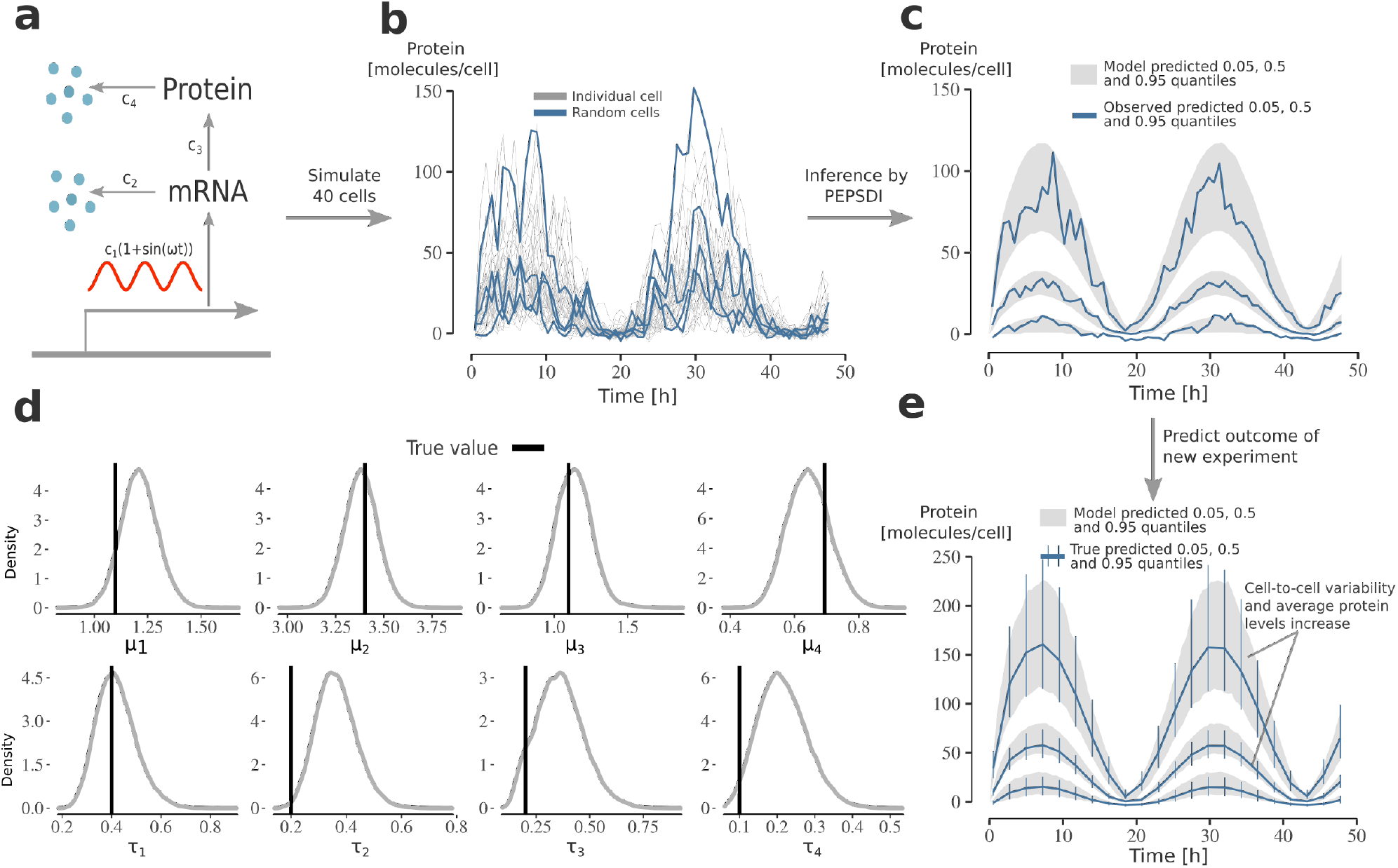
Applying PEPSDI on synthetic data from circadian clock gene expression model. **a)** Schematic representation of the gene expression model. The model consists of two states (mRNA, Protein), and four reactions with associated rate constants **c** = (*c*_1_, . . . , *c*_4_). The circadian clock, modelled as a sine function with a period of 24 hours, regulates the transcriptions activity *c*_1_. Extrinsic noise was simulated by assuming that rate-constants jointly follow a log-normal distribution: 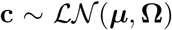. **b)** Protein count per cell for 40 cells simulated using the gene-network model. Data was simulated using the exact Extrande algorithm with an additive Gaussian measurement noise. **c)** Posterior visual checks [40]. The plot was generated as follows: i) from the inferred posterior distribution simulate 40 cells, and ii) for the 40 cells compute the 0.05, 0.5, 0.95 quantiles. This was repeated 10, 000 times yielding 95% credibility intervals. The blue lines correspond to the observed quantiles (from the data in (b)). **d)** Inference results using the data in b). The plots shows the marginal posterior for ***μ*** = (*μ*_1_, . . . , *μ*_4_) and ***τ*** = diag{**Ω**}^1/2^. **e)** Using the inferred model to predict the outcome of adding an extra gene in the model. An extra gene was modelled by doubling the transcription rate *c*_1_. The grey lines represent 95% credibility intervals for the 0.05, 0.5 and 0.95 quantiles (obtained as in subplot (c)) when simulating the inferred model with *c*_1_ doubled. The blue lines and error bars represents the true 95 % credibility intervals when observing 40 cells.

Additional extrinsic noise was simulated by letting the cell-specific rate constants follow a multivariate log-normal distribution; 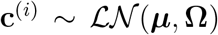 where ***μ*** and **Ω** are the mean and covariance matrix of the associated Gaussian distribution. Its multiplicative nature [26] makes the log-normal distribution a common choice for modelling parameters believed to vary between cells [6]. We also correlated the rate parameters to emulate that protein synthesis and degradation rates might co-vary [5, 27] (Supplementary Note 5).

The intrinsic noise was modelled using the exact Extrande simulator [15]. Even though faster approximate algorithms can be used to model intrinsic noise, low numbers of molecules in the model make these impractical [13].

We ran PEPSDI for 50, 000 iterations, and recovered the true model parameters of transcription rate, and how it varies within the cell population (Fig. 2d and Supplementary Fig. S1). There is a slight bias in the cell-to-cell variability of the breakdown rates (*τ*_2_, *τ*_4_), but, since the model accurately describes the data (Fig. 2c), this is likely because only 40 cells are observed.

The inferred model has predictive power. We considered an experiment where an additional gene is inserted into the model, and time-lapse data is collected for 40 new cells. The model accurately predicts an increase in both protein, and cell-to-cell variability levels (Fig. 2e).

### Application to simulated stochastic bistable model

Intrinsic noise can have a strong impact on cellular dynamics [28, 29]. Cellular processes have been shown to exhibit stochastic oscillations in gene regulation [28] and stochastic bistability as reported for the *lac*-operon regulation [29]. To study the performance of PEPSDI for such a process, we implemented the Schlögl model (Fig. 3a) [30], where cells stochastically migrate between states of high and low gene expression (Fig. 3b).

**Figure 3:**
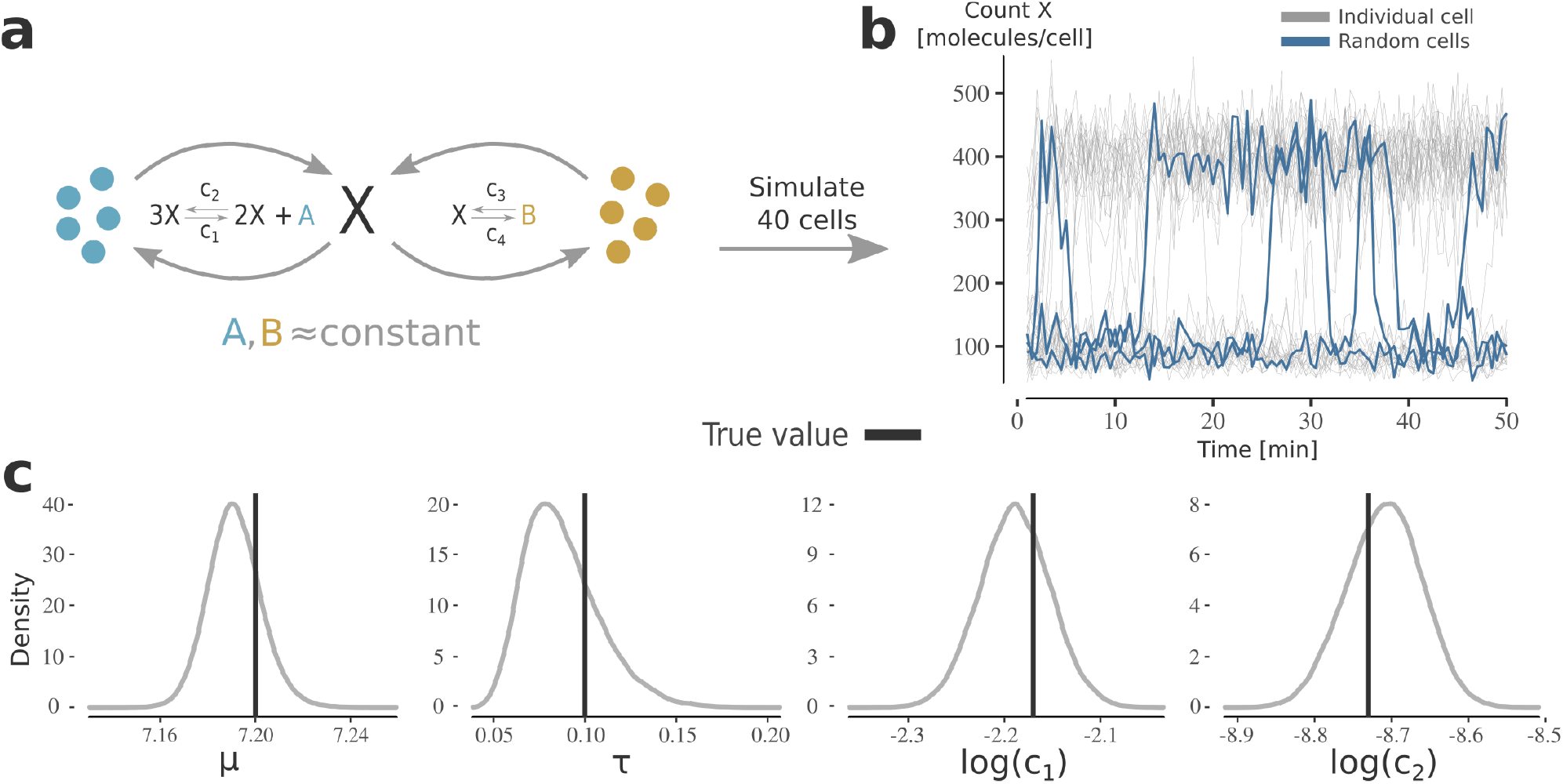
Applying PEPSDI on synthetic data from stochastic bi-stable model. **a)** Schematic representation of the stochastic bi-stable Schlögl model. Since the species (*A*, *B*) are assumed to be available in excess, the model consists of one state (X) and four reactions with associated rate constants **c** = (*c*_1_, . . . , *c*_4_). Extrinsic noise was simulated by assuming that *c*_1_ follows a log-normal distribution; 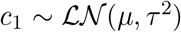. The remaining parameters are assumed to be cell-constant and *c*_4_ is assumed to be known. **b)** Molecule count of *X* per cell for 40 cells simulated using the Schlögl model. Data was simulated using the SSA algorithm with an additive Gaussian measurement noise. Noticeably, a subset of cells stochastically migrates between two cell-states. **c)** Inference results using the data in (b)). To efficiently infer the posterior distribution the model was simulated using the Langevin simulator. The plots shows the marginal posterior for the population parameters (*μ*, *τ*), and the cell-constant parameters (*c*_2_, *c*_3_).

To simulate extrinsic noise, we let one of the synthesis rates, *c*_3_, follow a log-normal distribution. To emulate that certain model parameters, such as protein dissociation rates, can have a neglectable variability [4], we kept a synthesis (*c*_1_) and dissociation rate (*c*_2_) constant between individuals. The synthesis rate *c*_4_ was assumed to be known (Supplementary Note 5).

To account for large numbers of molecules, we modelled intrinsic noise via the fast, approximate, Langevin simulator. We then used so-called guided proposals [9], directing simulations towards observed values, making the inference more efficient for models with stochastic events.

We ran PEPSDI for 500, 000 iterations, and posterior distributions of the modelled-quantities were recovered (Fig. 3c), allowing for better understanding of mechanistic properties of the model. For example, by simulating the inferred model, it becomes apparent that low values of the synthesis rate *c*_1_ commit cells to a first cell-state, e.g of low gene-expression, while for large values of *c*_1_, cells commit to, and jump between, two different states of gene expression (Supplementary Fig. S2).

### Performance evaluation of adaptive MCMC proposals

To propose unknown model quantities that rely on a pseudo-marginal step in our Gibbs sampler (see Methods), PEPSDI employs adaptive Markov chain Monte Carlo (MCMC) schemes [23]. However, these schemes were developed for problems where the likelihood function is not approximated via Monte Carlo scheme, as is the case for pseudo-marginal methods. Thus, we benchmarked three adaptive MCMC proposal schemes, namely adaptive metropolis (AM) [22], the AM with global scaling (Alg. 4 in [23]), and the robust AM (RAM) samplers [24]. For computational reasons, single time-series inference for two different models is considered.

For the Schlögl model (Fig. 3), the stochastic bistability (Fig. 3b) causes the likelihood approximation used in the pseudo-marginal method to have a large variance despite the usage of guided proposals. To investigate if this impacts the performance of adaptive MCMC schemes, we launched multiple inference runs. Overall, the RAM sampler on average had the highest multiple effective sample size (MultiESS) value [31], and thus most efficiently explored the posterior surface (Fig. 4a). This sampler also had the smallest variability in MultiESS, suggesting it is robust against, for example, bad start guesses (start guess 3 and 4 in Fig. 4a). Moreover, the RAM sampler typically required fewer particles, implying higher computational efficiency.

**Figure 4:**
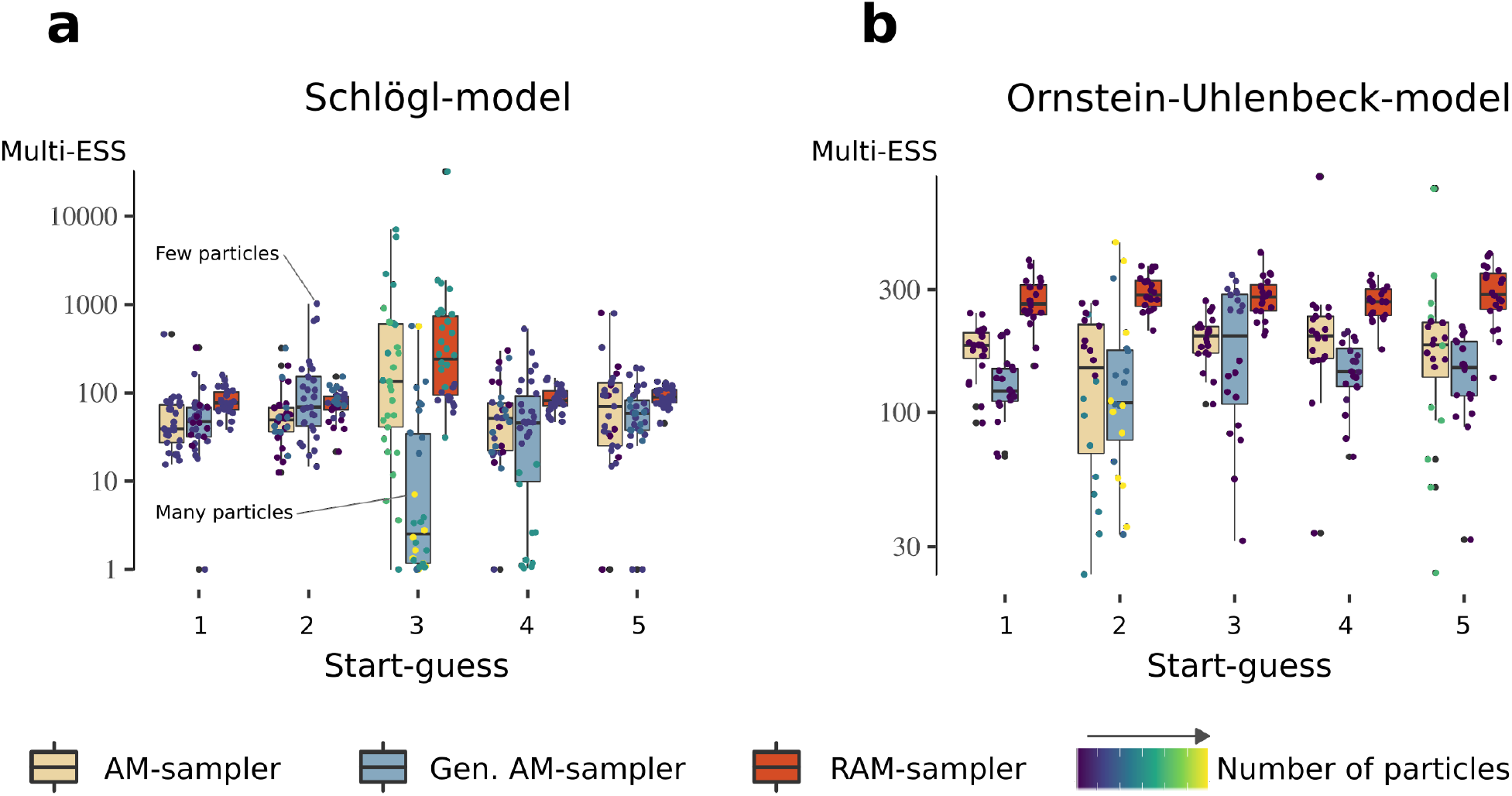
Benchmarking adaptive MCMC-proposals for pseudo-marginal inference. **a)** Benchmark results for the Schlögl-model. The quality of the adaptive schemes was measured using the MultiESS-criterium [31], where higher values are better. For computational reasons the benchmark was performed for a single-individual (single time-series data). Overall we simulated three datasets. For each dataset we ran five pilot runs with initial parameter values set at randomly chosen prior locations, and then tuned the number of particles (Supplementary Note 1). Starting from the last drawn parameter value in each pilot-run 10 further inference runs were independently launched. The colours denote the several adaptive proposal schemes, and the colour bar represents the number of particles selected by the tuning scheme. A high number of particles implies longer run-times, and an inefficient pilot run. **b)** Results for the Ornstein-Uhlenbeck model. The benchmark conditions were the same as in a).

For the Ornstein-Uhlenbeck stochastic differential equation model (Supplementary Note S5), the likelihood approximation has a small variance. Using the same setup as for the Schlögl model, the RAM sampler in this setting also performed the best.

### Application to glucose repression pathway

After considering synthetic examples, we set out to study the SNF1 pathway in *S.cerevisiae*. The SNF1 complex, and its mammalian homolog AMPK, play a major role in both metabolic regulation and maintenance of cellular homeostasis. In response to stress, such as ageing and nutrients limitation, SNF1 mediates the signal transduction to transcription factors. Mig1 is a transcriptional repressor which the SNF1 complex deactivates when energy-rich carbon sources are limited. This is followed by Mig1 relocalisation to the cytoplasm and release of repression of genes responsible for utilisation of alternative carbon sources [32, 33]. If the amount of energy-rich nutrients is elevated, Mig1 translocates to the nucleus [34]. This process is accompanied by Mig1 dephosphorylation where the Reg1 phosphatase plays the main role [1]. While this signalling cascade is well studied, it has also been indicated that the SNF1 pathway dynamics exhibits a high range of cell-to-cell variability [35].

To deduce the dynamics of the SNF1 pathway at a cell level, time-resolved data is required. We utilised fluorescent microscopy to follow Mig1 on a single-cell level and observe its localisation over time (Fig. 5a-b). This was coupled with microfluidics systems enabling high control of the cell environment. Mig1 localisation was followed during 15 min after cells were switched from carbon sourced depleted conditions to two different fructose conditions (w/v): 0.05%, and 2% (Fig. 5c).

**Figure 5:**
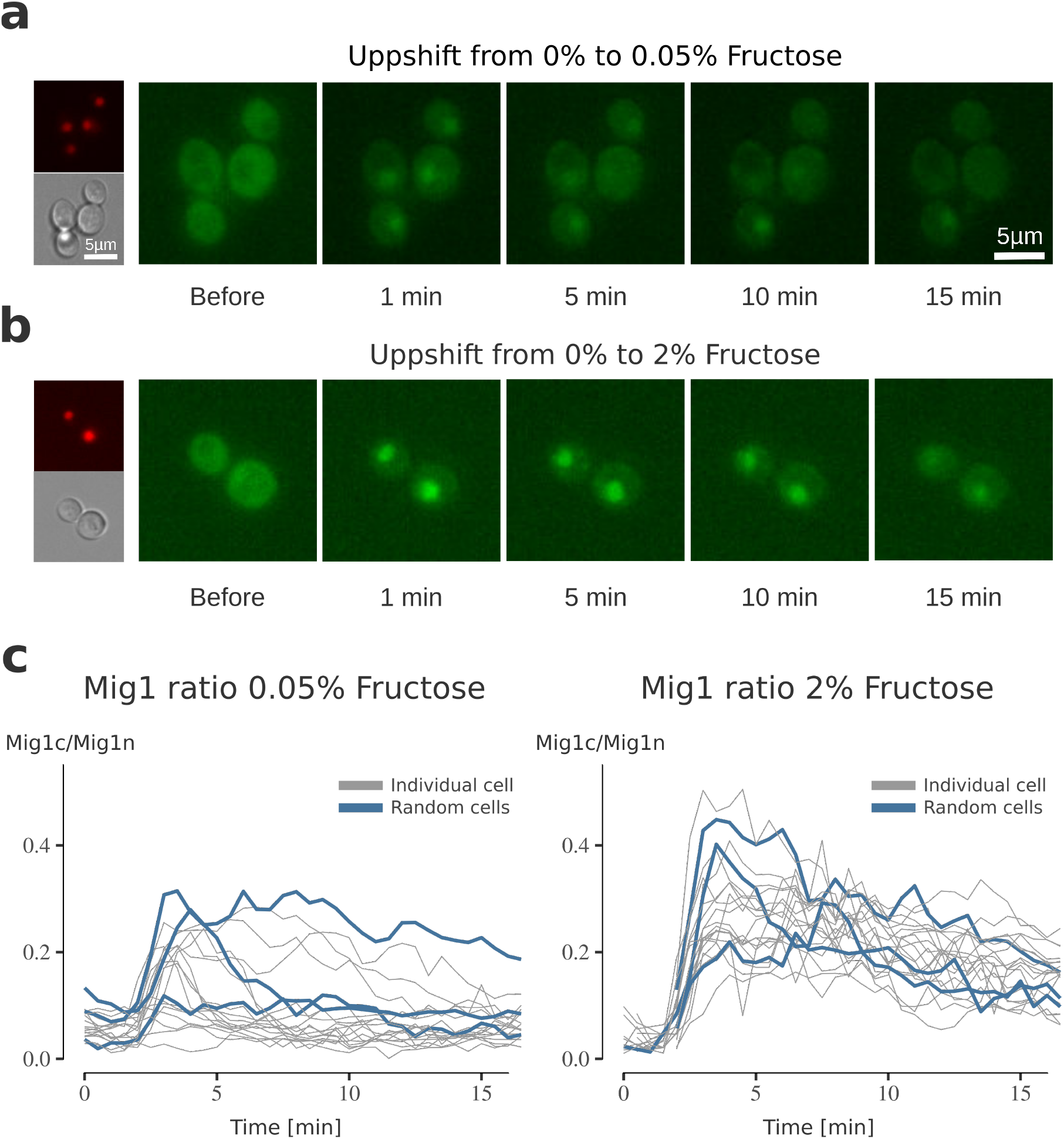
The impact of fructose on Mig1 dynamics. Mig1 localisation in response to the media exchange from media containing 0% fructose (no fast-fermentable carbon sources present) to media containing **a)** 0.05 % fructose and **b)** 2% fructose. The white lines corresponds to 5*μm* scale bars. Exchange of media was achieved through an open microfluidic system. Green fluorescent protein (GFP) images depict Mig1 localisation before and after switching of the media at the noted times. Brightfield images taken as control for the cell localisation. Red fluorescent protein (RFP) images depict Nrd1, a protein which is stationary in the nucleus and used as nuclear marker. **c)** The nuclear intensity of Mig1 for each single-cell in the experiment is given by the localisation index of Mig1 over time (minutes). Localisation index is determined by (Mig1n-Mig1c)/Mig1c (for short called Mig1n / Mig1c in the paper) with Mig1n being the intensity of Mig1 in the nucleus and Mig1c the intensity in the complete cell. All cell traces are grey, three random selected cells are given in blue. Overall, the data-sets combined consist of 37 individuals.

### Modelling of Mig1 nuclear dynamics

To elucidate both the source of cell heterogeneity and reaction mechanisms behind the Mig1 dynamics data (Fig. 5c), we formulated and analysed two network structures (Fig. 6a and b) and two extrinsic noise sources (Fig. 6c and d), resulting in four plausible models. In both network structures, Mig1 shuttles between the cytosol and nucleus in a carbon source-dependent [36] and -independent manner [37]. The carbon source-dependent response has been observed to occur in two phases, a transient initial Mig1 nuclear entry followed by nucleocytoplasmic shuttling [38]. This can be explained by two independent pathways regulating glucose derepression. One pathway activating the Snf1 kinase and the other pathway directing Snf1 towards Mig1 [39]. We investigated whether the first pathway promotes Mig1 nuclear entry via a fast signal (Fig. 6a), or if Mig1 nuclear entry is delayed due to Reg1 activation (Fig. 6b). The second pathway was modelled via an unknown metabolic component due to the fact that Mig1 dynamics are closely intertwined with metabolic activity [35, 38] (Supplementary Note 6).

**Figure 6:**
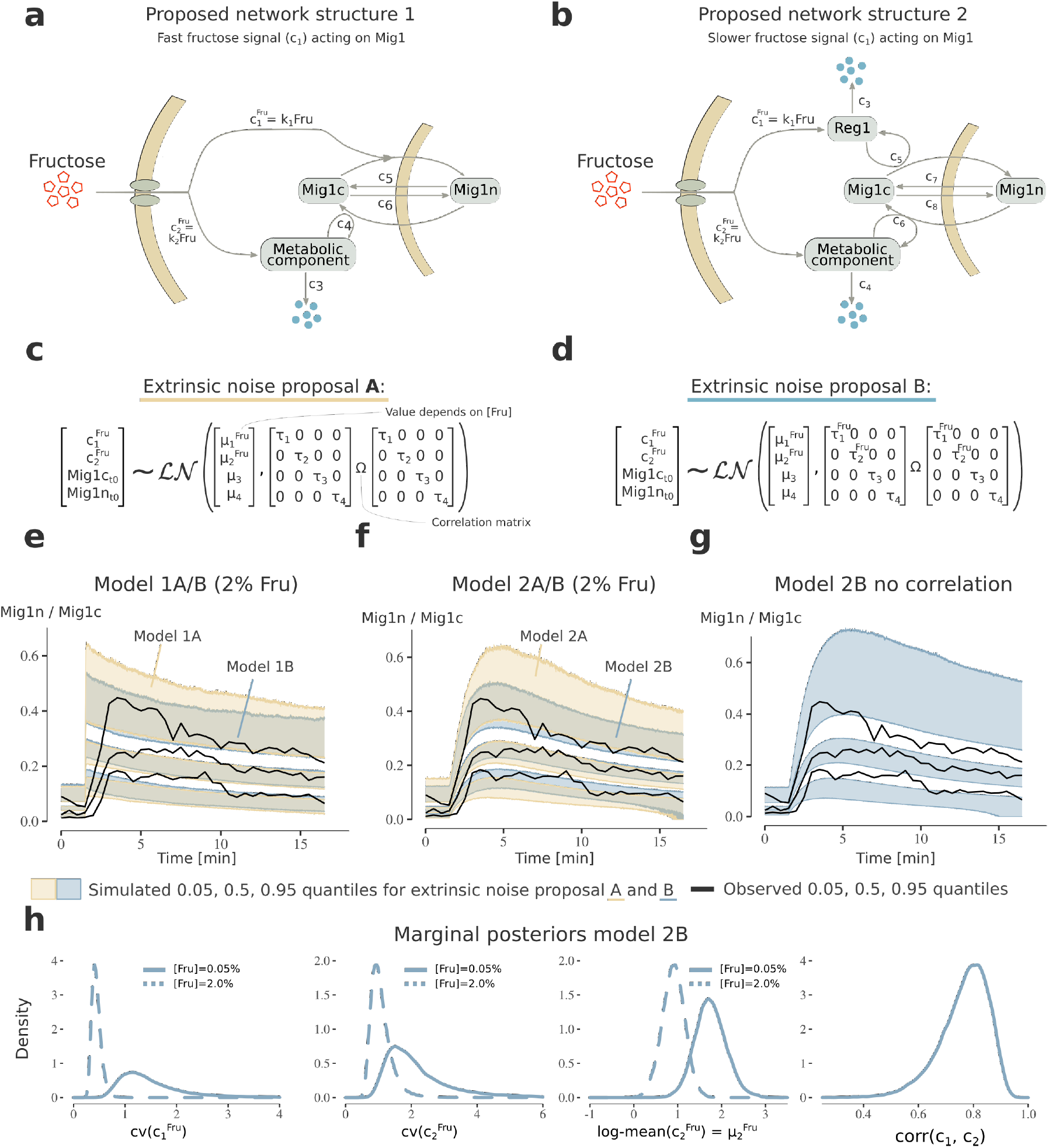
Modelling Mig1 dynamics in response to fructose addition. **a)** Proposed network structure 1 of Mig1 localisation dynamics. The model consists of three states, nuclear Mig1 (Mig1n), cytosolic Mig1 (Mig1c) and a metabolic component. The model proposes that a fructose signal directs Mig1 nuclear import, and that nuclear export is directed by a metabolic signal. **b)** Proposed network structure 2. In addition to structure 1 Mig1 nuclear entry is modelled as delayed due to Reg1 activation. **c)** Modelling proposal A of extrinsic noise. Since (*c*_1_, *c*_2_) encompass multiple cellular processes, and initial Mig1-levels are cell-varying, extrinsic noise was modelled by letting these follow a full log-normal distribution. Since (*c*_1_, *c*_2_) include the external fructose we model different log-mean values for the 2% and 0.05% conditions. **d)** Extrinsic noise proposal B. Additionally to proposal A, we model fructose dependent variability (*τ* -values) for (*c*_1_, *c*_2_). **e-f)** Posterior visual check for the 2% fructose data for model structure 1 and 2, using extrinsic noise-proposal A and B. The credibility intervals (bands) were obtained as in Fig. 2. Black lines are the observed quantiles. Model structure 2 with extrinsic noise proposal B (model 2B) best describes the observed trend, and cell heterogeneity. **g)** Posterior visual check as in f) for model 2B with no correlation between (*c*_1_, *c*_2_, *Mig*1*n, Mig*1*c*) (diagonal **Ω**). This yields an increase in cell-to-cell variability seen by the 0.05, and 0.95 quantile credibility intervals. **h**) Marginal posterior distribution for a subset of parameters in model 2B. The log-normal coefficient of variations 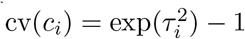 are fructose dependent, and (*c*_1_, *c*_2_) are strongly correlated.

To model the extrinsic noise sources, we assume that the Mig1 initial values (*Mig*1*c*_*t*0_, *Mig*1*n*_*t*0_) and the strength of the Mig1 regulation (*c*_1_, *c*_2_) vary between cells (Fig. 6c), as the latter incorporates multiple processes in the initial glycolysis. Since the rates (*c*_1_, *c*_2_) also encompass fructose abundance we assume and infer fructose-dependent log-mean values. The second extrinsic noise source proposal (Fig. 6d) takes into account that the cell-to-cell variability might be fructose-dependent.

We employed literature supported priors (Supplementary Note 6) and ran PEPSDI multiple times for each proposed model. Models were compared using posterior visuals check [40], that is the capability to capture both the observed trend and cell-to-cell variability (Fig. 6e, f and Supplementary Fig. S4). Overall, the model with a delayed fructose activation of Mig1 nuclear export and a fructose-dependent cell-to-cell variability in Mig1 regulation best described the data (model 2B Fig. 6f).

### Cell-to-cell variability in Mig1 nuclear dynamics is tightly regulated by fructose availability

After selecting the best model, we investigated the characteristics of the inferred model parameters. The coefficient of variations for the Mig1 regulation (*c*_1_, *c*_2_) shows that these rates substantially differ between cells (Fig. 6h). This suggests that extrinsic noise has a dominating role in Mig1 localisation dynamics. Furthermore, the magnitude of the variation, particularly for rate of activation of Reg1 via fructose (*c*_1_), is larger for low fructose conditions.

Our results show a correlation between *c*_1_, *c*_2_ rates (Fig. 6h), implying that cells with a strong fructose-dependent nuclear import of Mig1, also have a stronger nuclear export. By simulating the model without correlation we confirm that this correlation regulates cell-to-cell variability (Fig. 6g). Thus, our result suggests that co-regulation of two fructose-dependent pathways controls cell heterogeneity in Mig1 localisation dynamics.

The log-mean value of the rate parameter activating the metabolic component (log-mean(*c*_2_)) shows that the magnitude of the long-term nuclear export of Mig1 is weaker in high fructose (Fig. 6h). This is consistent with previous reports that nuclear export of Mig1 is primarily an effect of sugar depletion [36].

### The hexokinase Hxk1 is a source of cell heterogeneity in Mig1 localisation dynamics

In model simulations, the cell-varying rate constant *c*_1_ linearly correlates with the short-term (0-15 min) Mig1 localisation upon fructose addition to sugar-starved cells (Supplementary Fig. S4). Thus, since *c*_1_ captures a signalling process acting on Reg1 from the initial hexose metabolism via the hexokinase Hxk1, our modelling suggests a relationship between Hxk1 and such cell-to-cell variability.

To validate this prediction, we collected single-cell time-lapse microscopy data from a *hxk1* Δ*hxk2*Δ strain carrying Hxk1-expressing plasmids where both Mig1 localisation and Hxk1 expression are monitored upon fructose addition. In line with model predictions, the observed Hxk1 expression linearly correlates with Mig1 localisation, and the magnitude of variability in localisation explained by Hxk1 (linear regression *R*^2^-value) matches the predicted magnitude (Supplementary Fig. S4). We hence conclude Hxk1 as a source of cell heterogeneity in Mig1 dynamics.

## Discussion

Understanding the inherited nature of how biological processes dynamically change over time, and exhibit intra- and inter-individual variability, has been a major focus of systems biology.

The rise of single-cell fluorescent microscopy has enabled the study of those phenomena, but further progress is limited by the availability of methods that facilitate modelling, the essential follow up for rationalisation of such data. To address this, we developed PEPSDI, a versatile framework for Bayesian inference for dynamic stochastic single-cell models. We used PEPSDI to recover true model quantities for a circadian clock stochastic gene expression model, and deduce mechanistic details for a model where cells stochastically move between two states of gene expression. We hereafter studied the SNF1 signalling in yeast and identified hexokinase activity as a source of extrinsic noise, and deduced that sugar availability dictates cell-to-cell variability.

Modelling Mig1 dynamics (Fig. 6) suggests larger cell heterogeneity in the fructose activation of Reg1 (*c*_1_) and the component that regulates Mig1 shuttling (*c*_2_) upon fructose limitation (Fig. 6h). We hypothesise the presence of a sugar-dependent biological switch, triggered when fructose is present. This is consistent with the fact that under low fructose, only a few cells due to extrinsic noise activate the switch leading to larger cell-to-cell variability.

Our modelling also suggests cell variability in the Mig1 nuclear export. In particular, in the rate constant *c*_2_ which activates the metabolic component regulating Mig1 nuclear export. Although this process is slower than the Reg1 activation (*c*_1_), they are strongly correlated. Moreover, results show that the magnitude of *c*_2_ is larger under fructose limitation (Fig. 6h). It has been reported that rapid activation of the SNF1 complex upon glucose starvation is ensured by increased levels of ADP [41]. We therefore suggest that within the SNF1 pathway, ADP is a central part of the *c*_2_ rate constant, facilitating nuclear export of Mig1 via activation of the SNF1 complex. This is consistent with elevated ADP levels upon limited glucose or fructose concentrations in the cellular environment. Moreover, hexokinases that we identified to be a part of the *c*_1_ rate, regulate Reg1, which in turn also requires the abundant presence of a hexose sugar for full activity of the phosphatase [42]. Taken together, we hypothesise that *c*_1_ and *c*_2_ incorporate hexokinases and ADP, respectively, and are both controlled via the metabolism. This creates a tight co-regulation of the SNF1 pathway, which regulates cell-to-cell variability (Fig. 6g).

To study the single-cell behaviour of the Mig1 dynamics, we leveraged on the modifiable modular nature of PEPSDI. This modularity facilitates modelling of intrinsic noise by either the SSA [11], Extrande [15], tau-leaping [43] or Langevin [12] stochastic simulators. Additionally, new modules such as the hybrid-simulators [44] used to study NF*κ*B-pathway [45] can be incorporated. Likewise, new pseudo-marginal modules can be added, and considering that these schemes are actively researched [46, 47], the performance of the framework can improve further.

Besides being modifiable, we aimed to make PEPSDI accessible, by providing extensive tutorial notebooks on both how to use it, and how to leverage the underlying pseudo-marginal framework to model single time-series data. Additionally, to help users of pseudo-marginal inference we evaluated adaptive Markov chain Monte Carlo (MCMC) proposals [22, 23, 24], resulting in the best performance of the RAM sampler [24] (Fig. 4). However, the evaluation is performed on two models, and further analysis is required before generalising our conclusions.

In summary, we developed and employed PEPSDI to deduce the reaction dynamics, and sources of cell heterogeneity behind single-cell time-lapse data. Since PEPSDI is an inference framework for dynamic state-space mixed-effects models, it can also be applied for problems arising in ecology [9], in neuroscience [10] and in pharmacokinetics and pharmacodynamics (PKPD) [48]. With the versatility of applications for state-space mixed-effects models, and with our framework that does not impose strict model assumptions, it is easy to envision additional applications such as modelling the behaviour of cancer cell populations [49]. We believe that this framework will play an increasingly important role in addressing challenging biological questions that cannot be answered by experimental approaches only, thus providing novel insights for better understanding life processes.

## Materials and methods

### Stochastic simulations

Under the assumption that the system is well mixed, the dynamics of a stochastic reaction-network of *d* species, **X** = (*X*_1_(*t*), . . . , *X*_*d*_(*t*)), is described by the chemical master equation [13, 43]. The propensity function *h*(**x**, **c**, *t*) is a measure of the probability that one, out of *R*, reactions will occur depending on **X** and the rate-constants **c**. Considering that the master-equation can rarely be solved due to its probabilistic nature [13, 43], our inference framework relies on simulating from it.

The SSA direct method [11] produces exact stochastic simulations. Similarly, the Extrande simulator [15] produces exact solutions when the rate-constants **c** are time variant. However, both approaches simulate each reaction event and can thus be slow for large propensities [50]. Assuming that the propensities do not change noticeably during the time interval [*t*, *t* + *τ*] (leap condition 1), the reactions will be close to independence of each other. Hence, the tau-leap approach with fixed step-length can be used to update the states-vector from time *t* to *t* + *τ* [12, 51]. Further assuming that the propensities are sufficiently large in [*t*, *t*+*τ*] (leap conditions 2), the state-vector can be updated via the chemical Langevin stochastic differential equation [12, 43]. Typically, the number of molecules must be large for leap condition 1 and 2 to hold.

Our framework allows models to use the SSA method and the Extrande method when neither leap conditions holds. When one or both conditions holds, our framework allows usage of either tau-leaping or the Langevin approach.

### PEPSDI: Bayesian inference for single-cell dynamic models

PEPSDI performs Bayesian inference for state-space models with latent dynamics incorporating mixed-effects, shortly “state-space mixed-effects models” (SSMEMs). A state-space model is a discrete-time, stochastic model that contains two sets of equations: (i) one describing how a latent Markov process transitions in time (the state equation) and (ii) another one describing how an observer measures the latent process at each discrete time-point (the observation equation), assuming conditional independence between observations and latent states. Thus, a state-space model can describe a stochastic chemical reaction network. Here, we outline PEPSDI, and a more complete description also showing pseudo-algorithms is provided in Supplementary Note S1.

To perform inference, PEPSDI requires single-cell time-lapse measurements 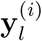 for the *i*-th individual collected at (*l* = 1, ..., *n*_*i*_) discrete time-points 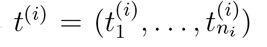 for *i* = 1, . . . , *M* individuals. Note, from now we denote with "individual" the measurements from a single cell. However since PEPSDI can be used in other applied areas (e.g. ecology, PKPD, etc), an individual is more generally a unit from the population of interest. The individual data is assumed to be noise-corrupted as 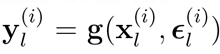, where we use the shorthand notation 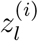 to denote a variable *z* observed at time 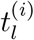. Here, 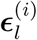 is the measurement error which follows an error distributio 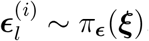 and **g**(·) s a (possibly non-linear) function of its arguments. To infer parameters in the nutrient-sensing Mig1 pathway, the observed Mig1 data represents a ratio of nuclear to cytosolic intensity, and thus 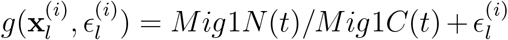, with independent 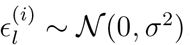.

For a SSMEM, PEPSDI infers the individual rate-constants **c** = (**c**^(1)^, . . . , **c**^(*M*)^), cell-constant rate-parameters ***κ***, the parameters ***ξ*** for the measurement error, and the population parameters ***η***. The population parameters describe the distribution of the individual parameters, **c**^(*i*)^ ~ *π*(**c**|***η***). For example if **c**^(*i*)^ follows a log-normal distribution, the population parameter corresponds to ***η*** = (***μ***, **Ω**) and individual rate constants to 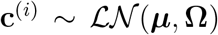. Overall, the posterior distribution we target is:

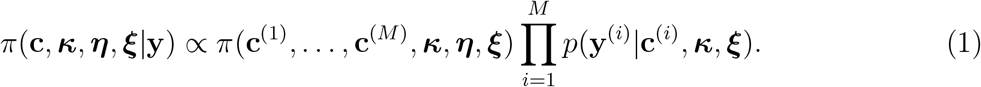

Note, in Eq .1 we have assumed that measurements from different subjects are conditionally independent, given the individual-specific **c**^(*i*)^ and the population parameters.

The posterior (Eq. 1) is high-dimensional, and ideally we could sample from it via a Gibbs-sampler [52] by looping through the following three steps:

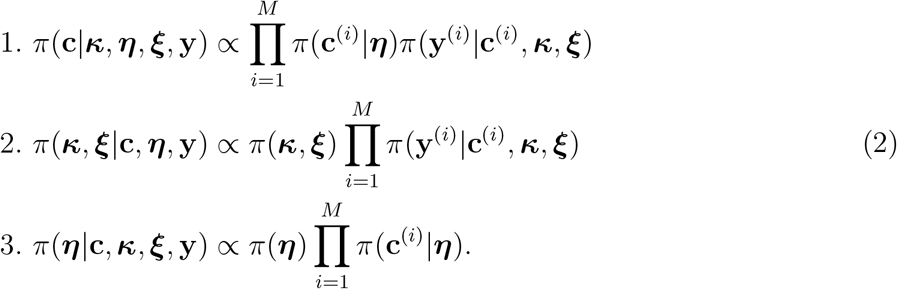

Notice that it is possible to sample from the first step by independently sampling for each of the **c**^(*i*)^ separately from the other ones. That is step 1 can be written as

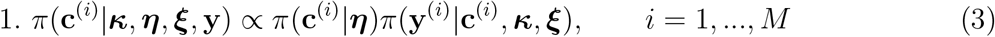

and hence the sampling step for each **c**^(*i*)^ only needs to access the corresponding individual-specific *π*(**y**^(*i*)^|**c**^(*i*)^, ***κ**, **ξ***).

However, in practice, steps 1-2 cannot trivially be sampled from, due to the intractability of the likelihood for the *i*-th individual *π*(**y**^(*i*)^|***κ***, ***ξ***, **c**^(*i*)^), and here sampling is performed using a pseudo-marginal approach following [10]. The posterior targeted in step 3 is tractable, and thus ***η*** is sampled using Hamiltonian Monte Carlo [17, 53, 54].

PEPSDI can be run with two flavours. Both sample the conditionals in Eq. 2 via, when required, pseudo-marginal approaches. However, the default option is to slightly perturb the SSMEM. This prevents the need of step 2 in the Gibbs-sampler, resulting in substantially shorter run-time. To properly motivate this perturbation, we first cover pseudo-marginal particles-based inference.

### Pseudo-Marginal particles-based inference

The pseudo-marginal Metropolis-Hastings scheme samples from the desired posterior by considering the marginal of an augmented posterior [16, 55]. More details are given in Supplementary Note 1. Briefly, the pseudo-marginal approach considers

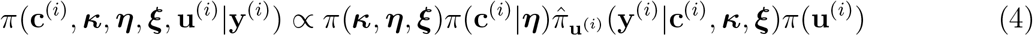

where often (though not necessarily) parameters can be a-priori independent *π*(***κ***, ***η***, ***ξ***) = *π*(***κ***)*π*(***η***)*π*(***ξ***), and the **u**^(*i*)^ ~ *π*(**u**^(*i*)^) are auxiliary variables used to obtain an unbiased estimate 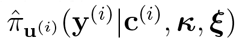 of *π*(**y**^(*i*)^|**c**^(*i*)^, ***κ***, ***ξ***), that is

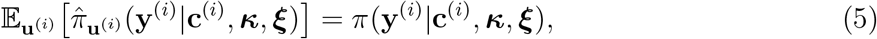

where 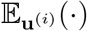 means that the expectation is taken with respect to the distribution of the **u**^(*i*)^. Thanks to the (assumed) unbiasedness of the estimated likelihood, the marginal of the augmented posterior is the *exact* posterior of interest

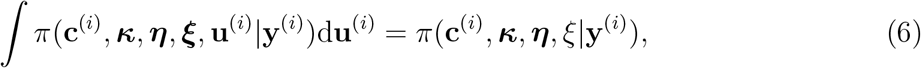

even though an estimated likelihood term has been employed inside Eq. 4. An efficient way to obtain an (non-negative) unbiased likelihood estimate for state-space models is to use a sequential Monte Carlo procedure known as the particle filter [56]. The particle filter approximates unbiasedly [57, 58] the expectation in Eq. 5 by using *N* Monte Carlo draws that in this context are named “particles”. The variance of the estimated likelihood decreases when increasing *N* , and typically the success of pseudo-marginal approaches relies on having a small likelihood-variance [47, 59, 60]. That is using too few particles yields inefficient inference, but on the other hand run-time increases with *N*. However, the result that makes pseudo-marginal powerful is that, from a theoretical point of view, it provides *exact* Bayesian inference [16, 55], regardless the number of particles employed, thanks to Eq. 6.

To employ as few particles as possible, the following three strategies are considered. Firstly, we induce correlation in the particles between subsequent iterations, and this still preserves exact Bayesian inference [21, 61]. However, this is only feasible for Poisson or Langevin integrators. Secondly, our framework implements a particles tuning scheme. Thirdly, guided particle proposals [9, 62] are used when possible (for details see Supplementary Note S1).

From the considerations above, PEPSDI performs exact Bayesian inference for the model parameters. However, to further reduce the computational requirements to run the inference scheme, we developed a Gibbs-sampler that targets a slightly perturbed parameterisation of a SSMEM-model, which we now describe.

### Inference for perturbed SSMEM (default option in PEPSDI)

The “perturbed SSMEM” treats cell-constant parameters (***κ***, ***ξ***) as parameters that instead vary, with a small fixed variance, between cells. For example, for ***ξ*** this means that for the perturbed SSMEM we assume to have 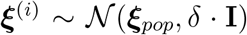, with *δ* > 0 a fixed constant selected by the researcher (and using a similar reasoning for ***κ***^(*i*)^). In some sense (***κ***, ***ξ***) have been “perturbed” to artificially vary between cells. This parameterisation, inspired by how cell-constant parameters can be treated in Monolix [63], allows (***ξ***_*pop*_, ***κ***_*pop*_) to be inferred alongside with the population parameters ***η*** in step 3, and (***ξ***^(*i*)^, ***κ***^(*i*)^) to be inferred alongside **c**^(*i*)^ in step 1. Step 2 is thus avoided.

Avoiding step 2 is desirable from the computational point of view. This is because step 2 of Gibbs-sampler (Eq. 2) requires a stochastic approximation of the sum of the individual log-likelihoods (for numerical stability it is preferable to work on log-transformed quantities), enabled by a particle filter. The variance of this sum is typically large, since each element of the sum is a stochastically approximated log-likelihood. To achieve a small-variance for this log-likelihood, and thus efficient inference, many particles are required which can cause substantial run-time (Supplementary Fig. S3).

As seen for the Ornstein-Uhlenbeck model, the Gibbs sampler corresponding to the perturbed SSMEM occasionally produces slightly wider credibility intervals for (***κ***_pop_, ***ξ***_pop_) (Supplementary Fig. S5). However, we consider this a worthwhile compromise since we do not observe any bias, while reaching a speed sometimes larger than a factor 30 (Supplementary Fig. S3). Moreover, if necessary PEPSDI can be run with the perturbed SSMEM to rapidly obtain preliminary results from pilot-runs, and the latter can be used to inform the setup (e.g. starting parameter values) to launch the inference for the unperturbed SSMEM.

### Single-cell microscopy data

A glass bottom petri-dish (GWST-5030, WillCo Wells) was treated with Poly-L-Lysine solution (P4832, Sigma-Aldrich) for 15 min at room temperature. The Poly-L-Lysine solution was removed, and the petri-dish was washed with MQ water for 2 times and left to dry overnight. Yeast cells (W303(202) NRD1-mCherry-Hph MIG1-GFP-KanMX) were grown overnight to mid-log phase at 30 °C in YNB synthetic complete medium (formedium) (6.7 g/l yeast nitrogen base with ammonium sulphate (formedium), 790 mg/l complete supplement mix (formedium) and supplied with 3% ethanol). These mid-log phase cells were added to the petri-dish and left to sediment, cell which did not adhere to the surface were removed by washing with growth media. Exposure of cells to YNB media with different concentrations of fructose was performed by using a BioPen system with BioPen prime pipette tip (Fluicell AB) as described in previous work [64]. Imaging was performed on an inverted microscope a Leica DMi8 inverted fluorescence microscope (Leica microsystems). The microscope was equipped with a HCX PL APO 40 × /1.30 oil objective (Leica microsystems), Lumencor SOLA SE (Lumencor) led light and Leica DFC9000 GT sCMOS camera (Leica microsystems). 3 datapoints were taken before the cells were exposed to fructose, after exposure a data point was taken every 30 second for 15 min. Every time point 5 images with an axial distance of 0.5 *μm* in between the images were acquired in transmission and fluorescent light to ensure an in-focus image for all cells. These images in transmission were taken with 10 ms exposure and 190 LED intensity. Mig1-GFP was observed with an filtercube with a excitation: 450/490, dichroic: 495 and emission: 500-550 filtercube at 300 ms exposure. Nrd1-mCherry was observed with an excitation: 540/580, dichroic: 585 and emission: 592-668 filtercube at 160 ms exposure at 30% LED intensity.

The Hexokinase experiment was performed in W3030(202) *hxk1* Δ*hxk2*Δ, Nrd1-mCherry, Mig1-GFP, TDH3p-HXK1-CYC1t, TDH3p-mAmetrine-CYC1t cells were exposed to an upshift in Fructose from 0 to 2.0%, experimental setup was as described in [38]. Analysis of fluorescence intensity was performed with the ImageJ distribution FIJI [65].

## Data availability

The .tif files of the the experimental data are available on figshare (https://figshare.com/s/2d56e0a6a928ef1dd7ac). The time-lapse data used to fit the Mig1-model are available on GitHub; https://github.com/cvijoviclab/PEPSDI/tree/main/Intermediate/Experimental_data/Data_fructose.

## Code availability

PEPSDI is available on GitHub (https://github.com/cvijoviclab/PEPSDI). The code for producing the results in this paper are further available in the GitHub repository (https://github.com/cvijoviclab/PEPSDI/tree/main/Code/Examples).

## Acknowledgements

This work was supported by the Swedish Research Council (VR2019-03924 to UP and VR2017-05117 to MC), the Chalmers AI Research Centre (CHAIR) to UP, the Swedish Foundation for Strategic Research (FFL15-0238 to MC) and the Marie Skłodowska-Curie grant agreement No 764591 to PR.

## Author contributions

S.P. and M.C. conceived the study. S.P. developed the method and Mig1 model, analysed and interpreted the data. S.S. and N.W. experimental design, interpretation of Mig1 data. N.W., P.R., G.S. carried out Mig1 experiments. S.W. assisted with method development, and U.P. co-supervised method development. M.C. supervised the work. S.P., N.W., S.S., U.P., M.C. wrote the paper, and all authors contributed to reviewing, editing and providing additional text for the manuscript.

