## Supplementary files for "PEPSDI: Scalable and flexible inference framework for stochastic dynamic single-cell models"

**S3 Guidelines for running our inference framework** **17**

**S4 A tutorial on constructing a single-cell dynamic model** **18**

**S5 Simulation examples** **20**

### S1 Supplementary figures

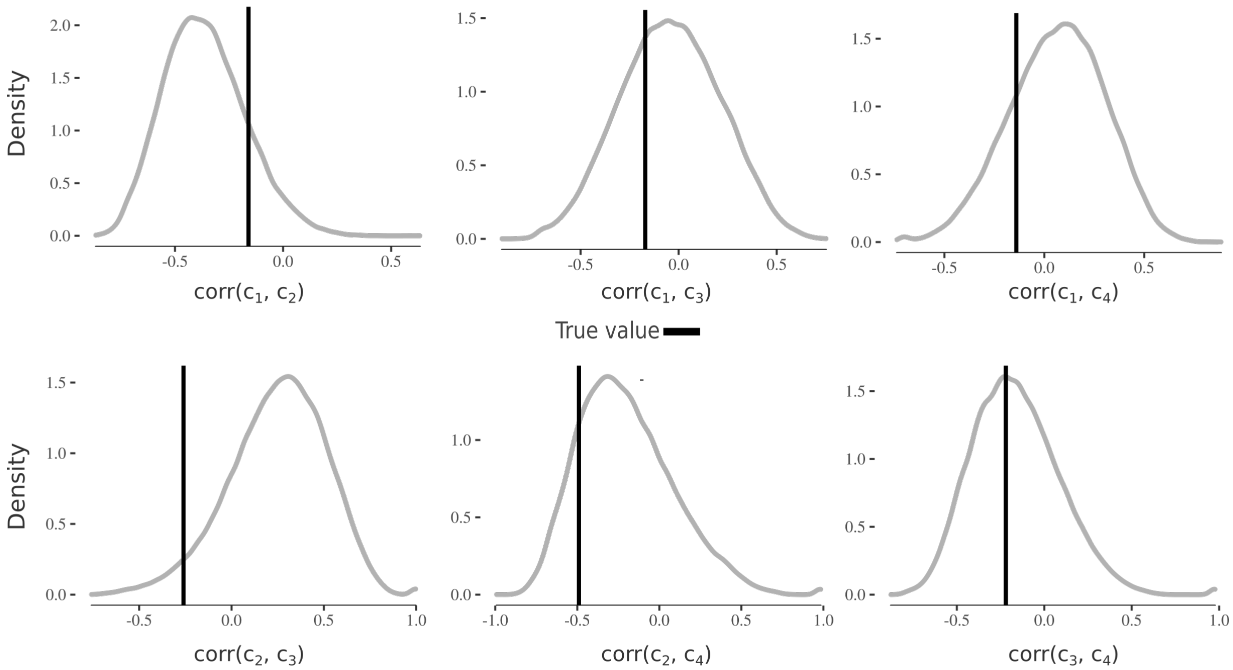

**Figure S1: Additional results for the stochastic gene-network regulated by the circadian-clock**. Marginal posterior from the inference run in Fig. 2 for the correlation matrix $\mathbf{\Phi}$ (non-diagonal values of the covariance matrix $\mathbf{\Omega}$). The correlation matrix characterises the correlation between the individual parameters $(c_1, c_2, c_3, c_4)$. The black line represents the true-value.

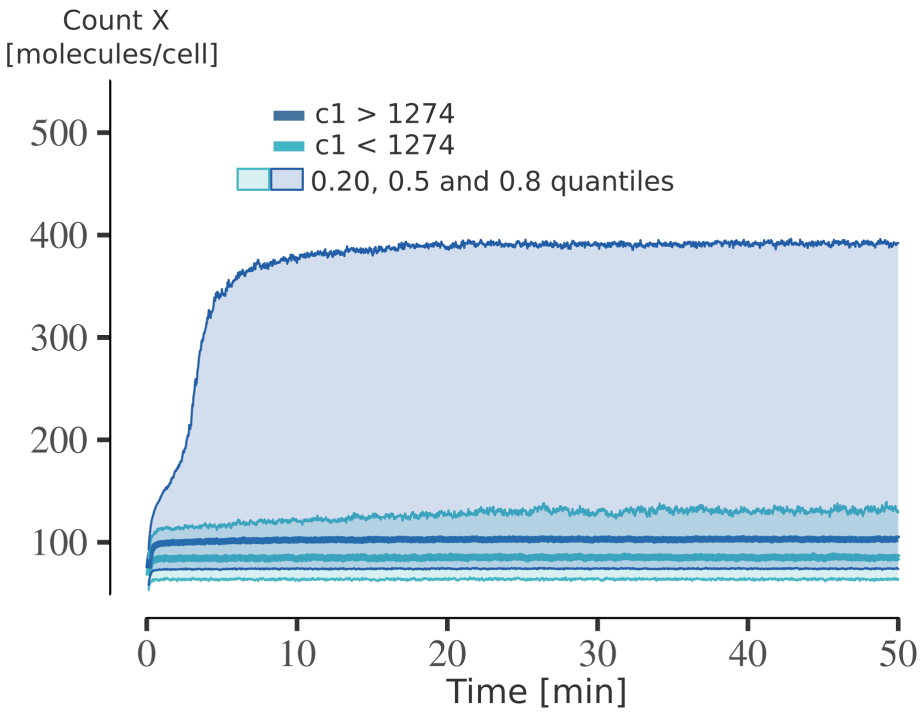

**Figure S2: Using inferred parameters to deduce mechanistic properties for the Schlögl-model.** Using the inferred posterior for the Schlögl-model (Fig. 3), $100,000$ cells were simulated. The cells were then split into the group having a synthesis rate $c_1$ below 1274, and above 1274. For these groups the 0.2, 0.5 and 0.8 quantiles were computed. As seen from these quantiles, cells with a lower synthesis rate ($c_1$) mainly commit to the lower cell-state (e.g low gene-expression). Meanwhile, cells with a larger synthesis rate commit to, and jump between, two different states of gene-expression.

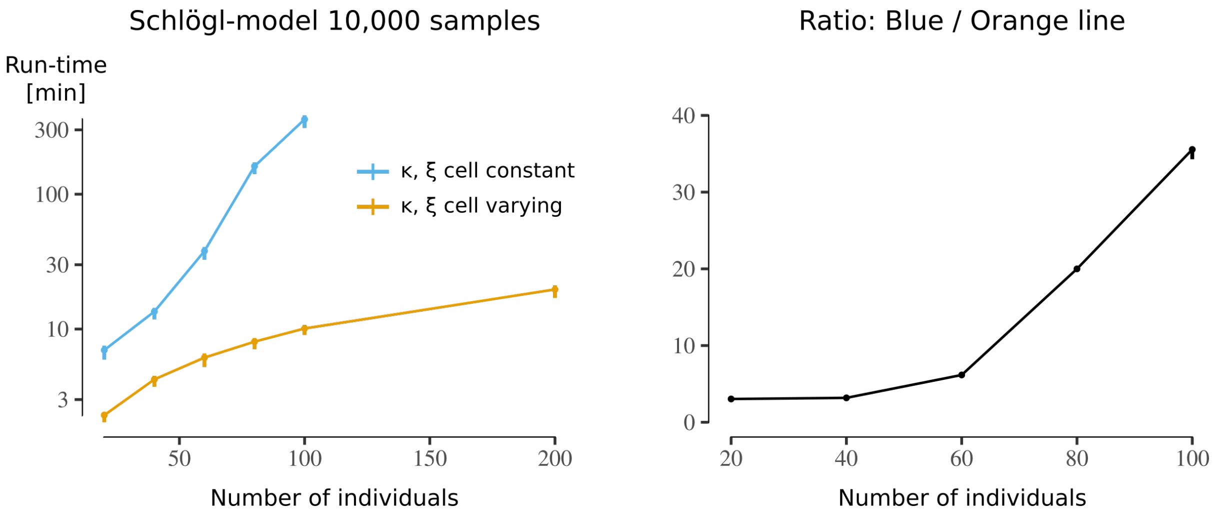

**Figure S3: Comparing run time of the PEPSDI inference options for the Schlögl-model.** **a)** Comparison of run-time for the non-perturbed option (blue) where $(\boldsymbol{\kappa}, \boldsymbol{\xi})$ are constant between cells, and the default perturbed option (orange), where $(\boldsymbol{\kappa}, \boldsymbol{\xi})$ are slightly perturbed to vary between cells. Using the same-parameters as in Fig. 3, data-sets for $20, 40, 60, 80, 100$ and $200$ individuals were simulated. Starting from true parameter values, the number of particles was tuned and PEPSDI was run for $10,000$ iterations. For all data-sets, the particle tuning procedure suggested the use of 10 particles per individual for the perturbed-model option (orange-line), while for the non-perturbed option (orange line) the procedure suggested to use (with increasing number of individuals) 20, 20, 40, 130 and 230 particles. The left plot shows the median run-time with max-and min values (bars) for three replicates. Due to computational burden run-time was not measured for the case of 200 individuals for the non-perturbed option. **b)** Ratio between the blue and orange line, highlighting that the default perturbed option can be faster by more than a factor 30.

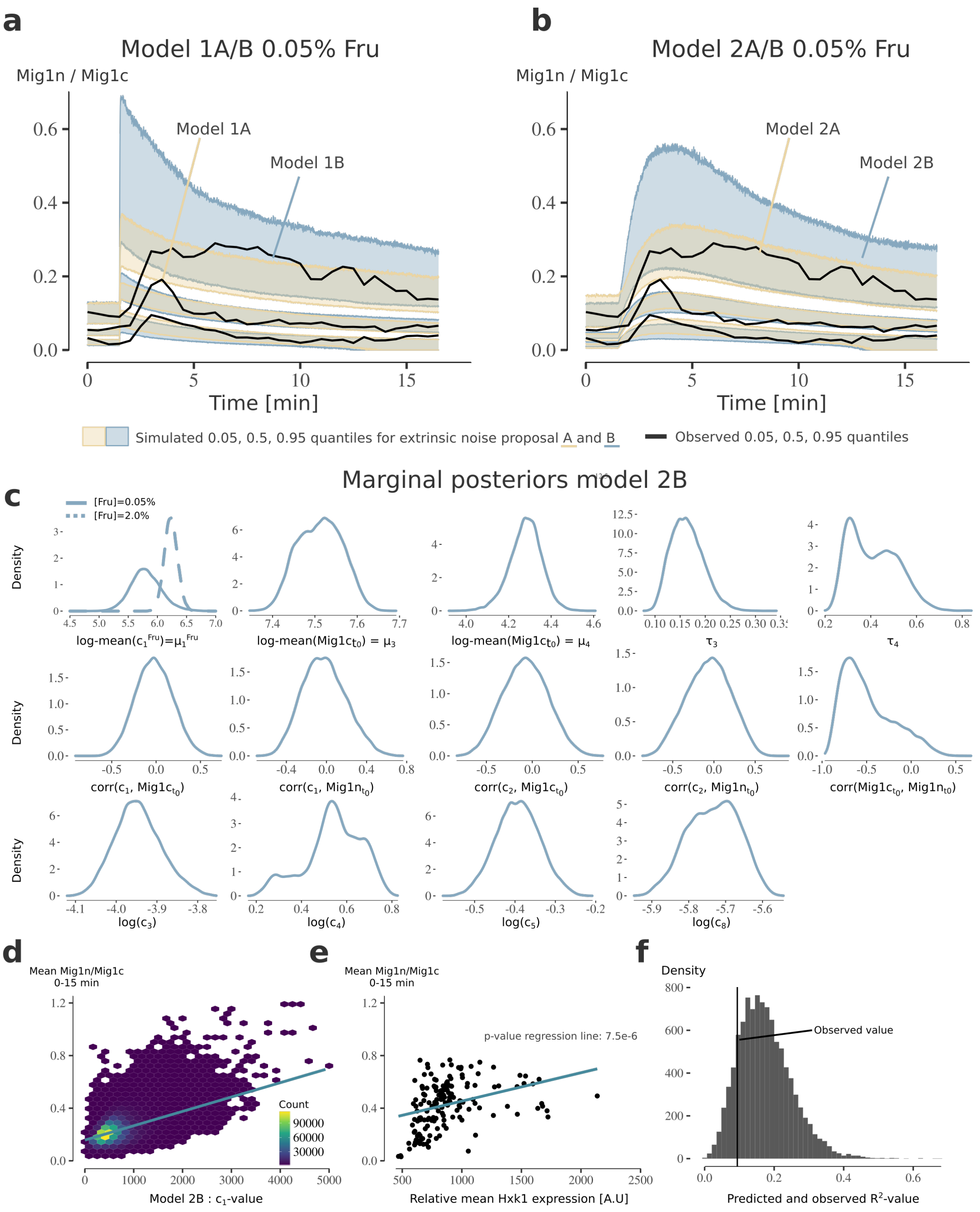

**Figure S4: Modelling of Mig1-dynamics in response to fructose addition. a-b)** Posterior visual check for the 0.05% fructose data for model structure 1 and 2 using extrinsic noise-proposals A and B (Fig. 6). The credibility intervals were obtained as in Fig. 2. Model 2B compared to model 2A has wider (but not biased) credibility intervals for the 0.05% fructose data, however, only model 2B accurately describes the 2% data (Fig. 6f). **c)** Marginal posterior for the model-parameters not shown in Fig. 6. **d**) Mean Mig1 ratio over 15 minutes after 2% fructose addition versus the cell-varying model parameters $c_1$ for model 2B obtained by simulating $1,320,000$ cells. Noticeable, the cell-varying $c_1$ of which hexokinase 1 (hxk1) is a part ($c_1 \propto [hxk1]$) explains a part of the cell-heterogeneity in Mig1 localisation. **e**) Mean Mig1 ratio over 15 minutes after 2% fructose addition versus relative hxk1 expression for 132 cells obtained from single-cell time-lapse microscopy. The linear relationship is significant (p-value $7.5 \times 10^{-6}$). The mean relative hxk1 expression, which is likely proportional against hxk1-expression, was computed as in [S1].

**Figure S4** ***(previous page)*****: f)** Model predicted (bars) and observed (line) explained cellular heterogeneity ($R^2$) in Mig1-localisation by relative hxk1-expression and $c_1$ respectively. The line is the $R^2$-value (variability explained by regression line divided by total variability) for the linear regression in e), and the bars were computed by simulating mean Mig1 ratio (0-15 minutes after fructose addition) for 132 cells $10,000$ times, and computing the $R^2$ for the Mig1n/Mig1c versus $c_1$ linear regression for each instance.

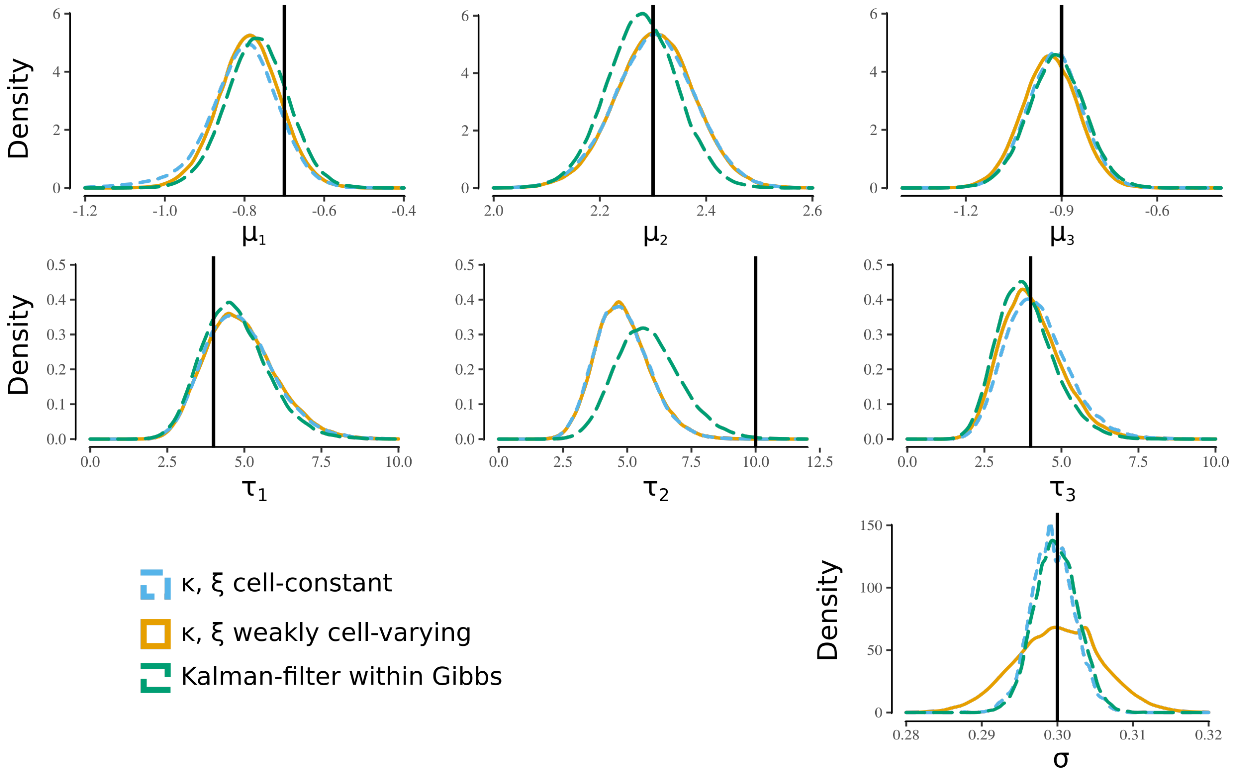

**Figure S5: Inference results for the Ornstein–Uhlenbeck model.** Inference was performed using PEPSDI with $(\boldsymbol{\kappa}, \boldsymbol{\xi})$ cell-constant and PEPSDI with $(\boldsymbol{\kappa}, \boldsymbol{\xi})$ weakly perturbed between cells (default option). These were compared against the gold-standard case, from Wiqvist et al. [S2], where a Kalman-filter is embedded into the Gibbs-sampler (Alg. 2) for an exact evaluation of the likelihood. Noticeably, our PEPSDI implementation performs well against the gold-standard case, suggesting a correct implementation. Furthermore, it can be seen that the consequence of perturbing the model is a slightly larger credibility interval for $\sigma$.

### S2 Inference framework

Our inference framework, PEPSDI, performs Bayesian-inference for state-space mixed-effects models (SSMEM) and was developed by extending the state-space mixed-effects inference framework for stochastic differential equations (SDEs) built in [S2] (see section S5.4 for a SDE example). We introduce two Gibbs samplers, with one named "modified Gibbs sampler", which is particularly efficient and is described in section S2.4. More precisely, for a SSMEM that describes a stochastic biochemical network of $d$-species, $\mathbf{X}^{(i)}(t, \mathbf{c}^{(i)}, \boldsymbol{\kappa}, \mathbf{D}^{(\mathbf{i})}) = (X_1^{(i)}, \ldots, X_d^{(i)})$ with $\mathbf{X}^{(i)} \in \mathbb{R}^d$, our framework infers unknown model quantities using observed single-cell time-lapse data $\mathbf{Y}^{(i)}(t_l^{(i)}) \in \mathbb{R}^{d_0}$, $l = 1, \ldots, n_i$, from $i = 1, \ldots, M$ individuals where $d_0 \leq d$. Here, it is assumed that the dynamics, $\mathbf{X}^{(i)}$ are governed by known cell-specific covariates $\mathbf{D}^{(i)} \in \mathbb{R}^m$, and unknown model-quantities. The unknown quantities are the individual rate-constants $\mathbf{c} = (\mathbf{c}^{(1)}, \ldots, \mathbf{c}^{(M)})$ where $\mathbf{c}^{(i)} \in \mathbb{R}^q$, the rate-constants with no assumed variability between individuals $\boldsymbol{\kappa} \in \mathbb{R}^p$, the parameters for the measurement error $\boldsymbol{\xi} \in \mathbb{R}^s$, and the population parameters $\boldsymbol{\eta} \in \mathbb{R}^r$ which parameterize the distribution of the individual parameters: $\mathbf{c}^{(i)} \sim \pi(\mathbf{c}^{(i)}|\boldsymbol{\eta})$. Here, it is always assumed that the data is noise-corrupted

$$\mathbf{y}(t_l)^{(i)} = \mathbf{y}_l^{(i)} = \mathbf{g}(\mathbf{x}_l^{(i)}, \mathbf{D}^{(i)}, \boldsymbol{\epsilon}_l), \quad \boldsymbol{\epsilon}_l \overset{iid}{\sim} \pi_{\boldsymbol{\epsilon}}(\boldsymbol{\xi}), \tag{S1}$$

where the $\boldsymbol{\epsilon}_l$ represents iid distributed measurement errors having distribution $\pi_{\boldsymbol{\epsilon}}(\boldsymbol{\xi})$ which is parameterised by a vector-parameter $\boldsymbol{\xi}$; $\mathbf{x}_l^{(i)}$ are the value of the model-states at time $t_l$ and $\mathbf{g}(\cdot)$ is a (possibly non-linear) function of its inputs. For the moment we do not add further restrictions to $\mathbf{g}(\cdot)$, however in some specific case we will need it to be a linear function of $\mathbf{x}_l^{(i)}$ and $\boldsymbol{\epsilon}_l$, see Eq. S15. Lastly, both $\boldsymbol{\kappa}$ and $\mathbf{c}^{(i)}$ can be augmented to include the initial system state values at time zero. As $\mathbf{D}^{(i)}$ equals the empty set throughout the paper, we hereafter discard $\mathbf{D}^{(i)}$ for simplicity of notation.

The unknown model quantities are inferred by sampling from the posterior

$$\pi(\mathbf{c}, \boldsymbol{\kappa}, \boldsymbol{\eta}, \boldsymbol{\xi}|\mathbf{y}) \propto \pi(\mathbf{c}^{(1)}, \ldots, \mathbf{c}^{(M)}, \boldsymbol{\kappa}, \boldsymbol{\eta}, \boldsymbol{\xi}) \prod_{i=1}^{M} \pi(\mathbf{y}^{(i)}|\mathbf{c}^{(i)}, \boldsymbol{\kappa}, \boldsymbol{\xi}), \tag{S2}$$

where $\mathbf{y} = (\mathbf{y}^{(1)}, ..., \mathbf{y}^{(M)})$ contains all observations across all $M$ individuals, and the likelihood term for the $i$-th individual can be written as

$$\pi(\mathbf{y}^{(i)}|\mathbf{c}^{(i)}, \boldsymbol{\kappa}, \boldsymbol{\xi}) = \prod_{l=1}^{n_i} \pi(\mathbf{y}_{t_l}^{(i)}|\mathbf{y}_{1:t_{l-1}}^{(i)}; \mathbf{c}^{(i)}, \boldsymbol{\kappa}, \boldsymbol{\xi}), \tag{S3}$$

and the notation $z_{1:l}$ means $(z_1, ..., z_l)$ for a generic variable $z$. For the posterior sampling we employ a Gibbs sampler. Some of the Gibbs-step have an intractable likelihood and for these we employ a pseudo-marginal approach as in [S2]. The tractable steps are sampled via efficient Hamiltonian Monte Carlo [S3, S4, S5]. Overall, this approach produces exact Bayesian inference, however, the run-time can be substantial. We thus have two options for running PEPSDI. The first option is the Gibbs approach in Alg. 2. The second (and the PEPSDI default option) is to run a Gibbs-sampler where cell-constant parameters are allowed to vary weakly between cells (Alg. 3). The latter considerably speeds-up the run-time (Fig. S3), with the potential caveat that the credibility intervals of the cell-constants parameters could be slightly wider (Fig. S5).

In this section we describe the inference framework in detail. Firstly, the different stochastic simulators currently implemented for simulating intrinsic noise are described. Secondly, the pseudo-marginal approach on which the Gibbs-samplers depend is described. Lastly, both our Gibbs-samplers are outlined.

#### S2.1 Stochastic simulators

For a well mixed system the dynamics of a biochemical stochastic reaction-network of $d$ species $\mathbf{X}^{(i)}(t, \mathbf{c}^{(i)}, \boldsymbol{\kappa}, \mathbf{D}^{(\mathbf{i})}) = \mathbf{X}_t^{(i)} = (X_1^{(i)}, \ldots, X_d^{(i)})$ with associated stoichiometry matrix $\mathbf{S} \in \mathbb{R}^{r \times d}$ is described by the chemical master equation [S6]. The key player here is the propensity vector $\mathbf{h}(\mathbf{x}_t^{(i)}, \mathbf{c}^{(i)}, t) \in \mathbb{R}^r$, where $h_j(\mathbf{x}_t^{(i)}, \mathbf{c}^{(i)}, \boldsymbol{\kappa}, t)\mathrm{d}t$ describes the probability that reaction $j$ will occur in the interval $[t, t + \mathrm{d}t)$. Typically, we cannot solve the master-equation so PEPSDI relies on simulations of its exact or approximate solutions. To accommodate a wide range of scenarios, our framework currently implements four different simulators.

The first simulator in our framework is Gillespie's "direct method" [S7]. This method produces exact stochastic simulations. However, the direct method assumes time-invariant rate-constants $\mathbf{c}$. To allow for time-varying rate-constants PEPSDI also accommodates the Extrande-simulator [S8]. Albeit exact, both the the Extrande- and Direct-method simulate each reaction event. Consequently, for large molecule numbers and/or large reaction rates, they can be slow [S6]. To reduce run-time our framework also allows for approximate simulators.

The third, and first approximate, simulator in our framework is the "tau-leaping" method with fixed step-length [S9, S10]. Assuming that the propensities do not change noticeably within $[t, t + \Delta\tau]$, that is assuming

$$h_j(\mathbf{x}^{(i)}, \mathbf{c}^{(i)}, \boldsymbol{\kappa}, t) \approx \text{constant} \quad \text{in } [t, t + \Delta\tau) \quad \forall j \qquad (1 : \text{st leap condition}), \tag{S4}$$

then the reactions will be close to independent one of the other. The reactions channels will thus fire independently, and the number $R_j$ of reactions for reaction channel $j$ is Poisson distributed as $R_j \sim \mathcal{P}\big(h_j(\mathbf{x}_t^{(i)}, \mathbf{c}^{(i)}, \boldsymbol{\kappa}, t)\tau\big)$. Overall, the state-vector can be updated via

$$\mathbf{x}_{t+\Delta\tau}^{(i)} = \mathbf{x}_t^{(i)} + \sum_{j=1}^{r} \mathrm{Row}_j(\mathbf{S})R_j, \quad R_j \sim \mathcal{P}\big(h_j(\mathbf{x}_t^{(i)}, \mathbf{c}^{(i)}, \boldsymbol{\kappa}, t)\Delta\tau\big). \tag{S5}$$

The fourth, and second approximate, simulator in our framework is the Langevin-chemical master equation. If, in addition to leap-condition 1, it is assumed that the propensities are sufficiently large, that is

$$h_j(\mathbf{x}_t^{(i)}, \mathbf{c}^{(i)}, \boldsymbol{\kappa}, t)\tau \gg 1, \ \forall j \qquad (2 : \text{nd leap condition}), \tag{S6}$$

then the Poisson-variable in Eq. S5 can be approximated by a Normal-distribution. Combining this with leap-condition 1, the state-vector can be described by a stochastic differential equation known as the Langevian chemical master equation [S9]:

$$\mathrm{d}\mathbf{X}_t^{(i)} = \underbrace{\mathbf{S}\mathbf{h}(\mathbf{X}_t^{(i)}, \mathbf{c}^{(i)}, \boldsymbol{\kappa}, t)}_{\boldsymbol{\alpha}(\mathbf{X}_t^{(i)}, \mathbf{c}^{(i)}, \boldsymbol{\kappa}, t)}\mathrm{d}t + \underbrace{\sqrt{\mathbf{S}\mathrm{diag}\big(\mathbf{h}(\mathbf{X}_\mathrm{t}^{(\mathrm{i})}, \mathbf{c}^{(\mathrm{i})}, \boldsymbol{\kappa}, \mathrm{t})\big)\mathbf{S}^T}}_{\sqrt{\boldsymbol{\beta}(\mathbf{X}_t^{(i)}, \mathbf{c}^{(i)}, \boldsymbol{\kappa}, t))}}\mathrm{d}W_t. \tag{S7}$$

Typically we cannot solve Eq. S7 analytically, so we use the Euler–Maruyama method to approximate the solution from time $t$ to $t + \Delta\tau$ as

$$\mathbf{x}_{t+\tau}^{(i)} \approx \mathbf{x}_t^{(i)} + \boldsymbol{\alpha}(\mathbf{x}_t^{(i)}, \mathbf{c}^{(i)}\boldsymbol{\kappa}, t)\Delta\tau + \sqrt{\beta(\mathbf{x}_t^{(i)}, \mathbf{c}^{(i)}, \boldsymbol{\kappa}, t)\Delta\tau} \cdot \mathbf{u}, \quad \mathbf{u} \sim \mathcal{N}(0, \mathbf{I}_{d \times d}), \tag{S8}$$

where $\mathbf{I}_{d\times d}$ is an identity unit matrix. As for the tau-leaping simulator an appropriate step-length $\Delta\tau$ must be chosen.

Overall, we implemented four simulators to allow the framework to handle a wide range of scenarios. When using the tau-leaping or the Langevin-simulators an appropriate step-length $\Delta\tau$ must be chosen. Approaches for doing this are given in S3. Owing to the modular nature of our code, it is straightforward to implement additional simulators such as hybrid-stochastic simulators [S11], or tau-leaping with an adaptive step-length [S12]. However, as highlighted in S2.5, it is beneficial for performance to use simulators where the number of pseudorandom numbers required for model simulation is known prior to starting the inference.

#### S2.2 Pseudo-marginal inference and particle filters

The joint posterior for individual rate-parameters $\mathbf{c} = (\mathbf{c}^{(1)}, \ldots, \mathbf{c}^{(M)})$, cell-constant rate-parameters $\boldsymbol{\kappa}$, population parameters $\boldsymbol{\eta}$, parameters for the measurement error $\boldsymbol{\xi}$, and latent data $\mathbf{x} = (\mathbf{x}^{(i)})_{i=1}^{M}$, given observed data $\mathbf{y} = (\mathbf{y}^{(i)})_{i=1}^{M}$, is

$$\pi(\mathbf{c}, \boldsymbol{\kappa}, \boldsymbol{\eta}, \boldsymbol{\xi}, \mathbf{x}|\mathbf{y}) \propto \pi(\mathbf{c})\pi(\boldsymbol{\kappa})\pi(\boldsymbol{\xi})\pi(\mathbf{c}|\boldsymbol{\eta})\pi(\mathbf{x}|\mathbf{c}, \boldsymbol{\kappa})\pi(\mathbf{y}|\mathbf{x}, \boldsymbol{\xi}),$$

where

$$\begin{aligned}
\pi(\mathbf{c}|\boldsymbol{\eta}) &= \prod_{i=1}^{M} \pi(\mathbf{c}^{(i)}|\boldsymbol{\eta}), \\
\pi(\mathbf{x}|\mathbf{c}, \boldsymbol{\kappa}) &= \prod_{i=1}^{M} \pi(\mathbf{x}_{t_1}^{(i)}) \prod_{l=2}^{n_i} \pi(\mathbf{x}_{t_l}^{(i)}|\mathbf{x}_{t_{l-1}}^{(i)}, \mathbf{c}^{(i)}, \boldsymbol{\kappa}), \\
\pi(\mathbf{y}|\mathbf{x}, \boldsymbol{\xi}) &= \prod_{i=1}^{M} \prod_{l=1}^{n_i} \pi(\mathbf{y}_{t_l}^{(i)}|\mathbf{x}_{t_l}^{(i)}, \boldsymbol{\xi}).
\end{aligned}$$

The posterior of interest is $\pi(\mathbf{c}, \boldsymbol{\kappa}, \boldsymbol{\eta}, \boldsymbol{\xi}|\mathbf{y})$, which is obtained via the following marginalisation

$$\pi(\mathbf{c}, \boldsymbol{\kappa}, \boldsymbol{\eta}, \boldsymbol{\xi}|\mathbf{y}) = \int \pi(\mathbf{c}, \boldsymbol{\kappa}, \boldsymbol{\eta}, \boldsymbol{\xi}, \mathbf{x}|\mathbf{y}) d\mathbf{x}. \tag{S9}$$

We thus have from (S9) that the parameter posterior is given by

$$\pi(\mathbf{c}, \boldsymbol{\kappa}, \boldsymbol{\eta}, \boldsymbol{\xi}|\mathbf{y}) \propto \pi(\mathbf{c})\pi(\boldsymbol{\kappa})\pi(\boldsymbol{\xi}) \prod_{i=1}^{M} \pi(\mathbf{c}^{(i)}|\boldsymbol{\eta})\pi(\mathbf{y}^{(i)}|\boldsymbol{c}^{(i)}, \boldsymbol{\kappa}, \boldsymbol{\xi}),$$

where $\pi(\mathbf{y}^{(i)}|\boldsymbol{c}^{(i)}, \boldsymbol{\kappa}, \boldsymbol{\xi})$ is the data likelihood for individual $i$. The data likelihood is obtained via the following marginalisation

$$\begin{aligned}
\pi(\mathbf{y}^{(i)}|\boldsymbol{c}^{(i)}, \boldsymbol{\kappa}, \boldsymbol{\xi}) &= \int \pi(\mathbf{y}^{(i)}, \mathbf{x}^{(i)}|\boldsymbol{c}^{(i)}, \boldsymbol{\kappa}, \boldsymbol{\xi}) d\mathbf{x}^{(i)}, \\
&\propto \int \prod_{l=1}^{n_i} \pi(\mathbf{y}_{t_l}^{(i)}|\mathbf{x}_{t_l}^{(i)}, \boldsymbol{\xi})\pi(\mathbf{x}_{t_1}^{(i)}) \prod_{l=2}^{n_i} \pi(\mathbf{x}_{t_l}^{(i)}|\mathbf{x}_{t_{l-1}}^{(i)}, \mathbf{c}^{(i)}, \boldsymbol{\kappa}) d\mathbf{x}_1^{(i)} d\mathbf{x}_2^{(i)} \cdots d\mathbf{x}_{n_i}^{(i)}.
\end{aligned} \tag{S10}$$

The integral in (S10) is generally intractable (except for simple cases, where the observations equation (S1) and the states dynamics are both linear in the states, and both have Gaussian noise, and in that case the Kalman filter can be applied) however we can use a pseudo-marginal approach when we replace the intractable integral with a unbiased estimation obtained via sequential Monte Carlo. Specifically, the pseudo-marginal approach allows for exact Bayesian inference when the observed data likelihood is intractable, but a non-negative and unbiased estimate is available. The pseudo-marginal Metropolis-Hastings scheme samples from the desired posterior via the marginal of an extended posterior [S13]. We have sketched some notions in the main paper, which we reprise here for convenience. In the pseudo-marginal approach we consider the posterior

$$\pi(\mathbf{c}^{(i)}, \boldsymbol{\kappa}, \boldsymbol{\eta}, \boldsymbol{\xi}, \mathbf{u}^{(i)}|\mathbf{y}^{(i)}) \propto \pi(\boldsymbol{\kappa}, \boldsymbol{\eta}, \boldsymbol{\xi})\pi(\mathbf{c}^{(i)}|\boldsymbol{\eta})\hat{\pi}_{\mathbf{u}^{(i)}}(\mathbf{y}^{(i)}|\mathbf{c}^{(i)}, \boldsymbol{\kappa}, \boldsymbol{\xi})\pi(\mathbf{u}^{(i)}) \tag{S11}$$

where often (though not necessarily) parameters can be a-priori independent $\pi(\boldsymbol{\kappa}, \boldsymbol{\eta}, \boldsymbol{\xi}) = \pi(\boldsymbol{\kappa})\pi(\boldsymbol{\eta})\pi(\boldsymbol{\xi})$, and the $\mathbf{u}^{(i)} \sim \pi(\mathbf{u}^{(i)})$ are auxiliary variables used to obtain an unbiased non-negative estimate $\hat{\pi}_{\mathbf{u}^{(i)}}(\mathbf{y}^{(i)}|\mathbf{c}^{(i)}, \boldsymbol{\kappa}, \boldsymbol{\xi})$ of $\pi(\mathbf{y}^{(i)}|\mathbf{c}^{(i)}, \boldsymbol{\kappa}, \boldsymbol{\xi})$, that is

$$\mathbb{E}_{\mathbf{u}^{(i)}}\left[\hat{\pi}_{\mathbf{u}^{(i)}}(\mathbf{y}^{(i)}|\mathbf{c}^{(i)}, \boldsymbol{\kappa}, \boldsymbol{\xi})\right] = \int \hat{\pi}_{\mathbf{u}^{(i)}}(\mathbf{y}^{(i)}|\mathbf{c}^{(i)}, \boldsymbol{\kappa}, \boldsymbol{\xi})\pi(\mathbf{u}^{(i)})d\mathbf{u}^{(i)} = \pi(\mathbf{y}^{(i)}|\mathbf{c}^{(i)}, \boldsymbol{\kappa}, \boldsymbol{\xi}). \tag{S12}$$

The assumed unbiasedness of the approximate likelihood ensures that the marginal of the targeted posterior (Eq.S11) is the desired (exact) posterior,

$$\int \pi(\mathbf{c}^{(i)}, \boldsymbol{\kappa}, \boldsymbol{\eta}, \boldsymbol{\xi}, \mathbf{u}^{(i)}|\mathbf{y}^{(i)})d\mathbf{u}^{(i)} \propto \pi(\boldsymbol{\kappa}, \boldsymbol{\eta}, \boldsymbol{\xi})\pi(\mathbf{c}^{(i)}|\boldsymbol{\eta}) \int \hat{\pi}(\mathbf{y}^{(i)}|\mathbf{c}^{(i)}, \boldsymbol{\xi}, \boldsymbol{\kappa}, \mathbf{u}^{(i)})\pi(\mathbf{u}^{(i)})d\mathbf{u}^{(i)} =$$
$$\pi(\boldsymbol{\kappa}, \boldsymbol{\eta}, \boldsymbol{\xi})\pi(\mathbf{c}^{(i)}|\boldsymbol{\eta}) \underbrace{\int \hat{\pi}_{\mathbf{u}^{(i)}}(\mathbf{y}^{(i)}|\mathbf{c}^{(i)}, \boldsymbol{\kappa}, \boldsymbol{\xi})\pi(\mathbf{u}^{(i)})d\mathbf{u}^{(i)}}_{=\pi(\mathbf{y}^{(i)}|\mathbf{c}^{(i)}, \boldsymbol{\kappa}, \boldsymbol{\xi})} \propto \pi(\mathbf{c}^{(i)}, \boldsymbol{\kappa}, \boldsymbol{\xi}, \boldsymbol{\eta}|\mathbf{y}^{(i)}). \tag{S13}$$

To obtain an unbiased [S14, S15] likelihood estimate we use a sequential Monte-Carlo procedure known as "particle filter" (Alg. 1), see [S16] for a recent review. Briefly, a particle filter recursively estimates a sequence of filtering densities $\pi(\mathbf{x}_{1:t}^{(i)}|\mathbf{y}_{1:t}^{(i)}, \mathbf{c}^{(i)}, \boldsymbol{\kappa}, \boldsymbol{\xi})$ for each $t_l, l = 1, \dots, n_i$, where $\mathbf{y}_{1:t}^{(i)}$ are the observations from time-step 1 to $t$ for the $i$-th individual. This is achieved by using a series of move and re-sampling steps. To move the particles a model dependent proposal distribution $q(\mathbf{x}_t^{(i)}|\mathbf{y}_t^{(i)}, \mathbf{x}_{t-1}^{(i)}, \mathbf{c}^{(i)}, \boldsymbol{\kappa}, \boldsymbol{\xi})$ is employed. This proposal distribution could correspond to a SSA-simulator, or an approximate simulator such as the tau-leaping simulator or the Euler-Maruyama method for SDEs, but the list could continue. Following a move, the particles are resampled (with replacement) to avoid so-called "particle impoverishment". A side-product of approximating the filtering distribution is that an unbiased non-negative estimate of the observed data likelihood is obtained via

$$\hat{\pi}_{\mathbf{u}^i}(\mathbf{y}^{(i)}|\mathbf{c}^{(i)}, \boldsymbol{\kappa}, \boldsymbol{\xi}, \mathbf{u}^{(i)}) = \hat{Z}^{(i)} = \frac{1}{N^{(i)}} \prod_{t=1}^{n_i} \sum_{k=1}^{N^{(i)}} \tilde{w}(\mathbf{u}_{t,k}^{(i)}), \tag{S14}$$

where $N^{(i)}$ is the number of particles used in the particle filter for the $i$-th individual. Less formally, the particle filter propagates forward in time multiple traces (particles) of the latent system's state from the assumed model. At each time step, the traces (particles) that are closer

**Algoritm 1** General particle filter

**Input**: Parameters $\mathbf{c}^{(i)}, \boldsymbol{\kappa}, \boldsymbol{\xi}$, data $\mathbf{y}^{(i)}$, auxiliary variables $\mathbf{u}^{(i)}$ and number of particles $N^{(i)}$.
**Output**: Observed likelihood estimate $\hat{Z}^{(i)}$ for individual $i$.
**Notation**: We use the convention that $k$ means "for all $k = 1, \dots, N^{(i)}$".

1: Initialisation ($t = 1$)

a: **Initialize the starting state**: set the value for $\mathbf{x}_{1,k}^{(i)}$ or sample it from some initial distribution $\mathbf{x}_{1,k}^{(i)} \sim q_1(\cdot)$.

b: **Compute the weights**:

$$\tilde{w}(\mathbf{u}_{1,k}^{(i)}) = \pi(\mathbf{y}_1^{(i)} | \mathbf{x}_{1,k}^{(i)}, \boldsymbol{\xi}), \quad w(\mathbf{u}_{1,k}^{(i)}) = \frac{\tilde{w}(\mathbf{u}_{1,k}^{(i)})}{\sum_{k=1}^{N^{(i)}} \tilde{w}(\mathbf{u}_{1,k}^{(i)})}$$

c: Compute the observed data likelihood estimate $\hat{Z}^{(i)} = \sum_{k=1}^{N^{(i)}} \tilde{w}(\mathbf{u}_{1,k}^{(i)})/N^{(i)}$

2: For $t = 2, 3, \dots, n_i$:

a: **(optional) Sorting**: if correlated particles are used, employ Euclidean sorting on $\{\mathbf{x}_{t-1,1}^{(i)}, \dots, \mathbf{x}_{t-1,N^{(i)}}^{(i)}\}$ to obtain sorted indices $s(k)$. Set $\{\mathbf{x}_{k,t-1}^{(i)}, w(\mathbf{u}_{k,t-1}^{(i)})\} := \{\mathbf{x}_{s(k),t-1}^{i}, w(\mathbf{u}_{s(k),t-1}^{(i)})\}$

b: **Resample**: sample (with replacement) $N^{(i)}$ particles from $\{\mathbf{x}_{t-1,1}^{(i)}, \dots, \mathbf{x}_{t-1,N^{(i)}}^{(i)}\}$ according to the normalised weights $w(\mathbf{u}_{k,t-1}^{i})$. Denote the resampled particles with $\{\tilde{\mathbf{x}}_{t-1,1}^{(i)}, \dots, \tilde{\mathbf{x}}_{t-1,N^{(i)}}^{(i)}\}$.

c: **Propagate** the particles $\mathbf{x}_{t,k}^{(i)} \sim q(\cdot | \mathbf{y}_t^{(i)}, \tilde{\mathbf{x}}_{t-1,k}^{(i)}, \mathbf{c}^{(i)}, \boldsymbol{\kappa}, \boldsymbol{\xi})$.

d: **Compute weights**:

$$\tilde{w}(\mathbf{u}_{t,k}^{(i)}) = \frac{\pi(\mathbf{y}_t^{(i)} | \mathbf{x}_{t,k}^{(i)}, \boldsymbol{\xi}) \pi(\mathbf{x}_{t,k}^{(i)} | \tilde{\mathbf{x}}_{t-1,k}^{(i)}, \mathbf{c}^{(i)}, \boldsymbol{\kappa})}{q(\mathbf{x}_{t,k}^{(i)} | \mathbf{y}_t, \tilde{\mathbf{x}}_{t-1,k}^{(i)}, \mathbf{c}^{(i)}, \boldsymbol{\kappa}, \boldsymbol{\xi})}, \quad w(\mathbf{u}_{t,k}^{(i)}) = \frac{\tilde{w}(\mathbf{u}_{t,k}^{(i)})}{\sum_{k=1}^{N^{(i)}} \tilde{w}(\mathbf{u}_{t,k}^{(i)})}$$

e: **Update the likelihood estimate**: $\hat{Z}^{(i)} = \hat{Z}^{(i)} \times \sum_{k=1}^{N^{(i)}} \tilde{w}(\mathbf{u}_{t,k}^{(i)})/N^{(i)}$

to observed data are favoured in the re-sampling step. Only the resampled particles are then propagated forward, and therefore the less promising ones "die".

There are multiple factors to consider when executing the particle filter. Firstly, any number of particles can be used. However, using too few increases the variance of the estimated likelihood, which has negative consequences for the mixing of the Markov-chain when this approximate likelihood with high variance is used in a Metropolis-Hastings step. An approach for selecting the number of particles is given in S2.5. Secondly, the particles can be moved using SSA-simulator, the tau-leap simulator, the Langevian-simulator or via guided proposals. Which propagator to use depends on the situation, and the propagators currently implemented in our framework are discussed in S2.2.1. Thirdly, the particles can be re-sampled in multiple ways. Following [S2] we use systematic re-sampling which has linear complexity. Notice, Alg. 1 contains an optional "sorting" step, which is only executed if a procedure meant at correlating the particles is employed, as motivated in S2.5.1.

##### S2.2.1 Propagators for particle filters

There are several ways to propagate particles forward in time, here we consider two approaches. The simplest is to propagate the particles forward in time using the latent state's transition density as proposal distribution, that is setting $q(\cdot|\mathbf{y}_t^{(i)}, \mathbf{x}_{t-1}^{(i)}, \mathbf{c}^{(i)}, \boldsymbol{\kappa}, \boldsymbol{\xi}) = \pi(\mathbf{x}_{t,k}^{(i)}|\mathbf{x}_{t-1,k}^{(i)}, \mathbf{c}^{(i)}, \boldsymbol{\kappa})$. This approach defines the so-called "bootstrap filter" [S17, S18], and it has evident appeal as it is often not clear how to construct a proposal distribution $q(\cdot)$ that depends on data, while it is always possible to propagate forward the system's state (or an approximation thereof), which essentially means simulating from the transition density (or an approximation thereof). Clearly, in the bootstrap filter the particles weights (step 2d Alg. 1) simplify to become $\tilde{w}(\boldsymbol{u}_{\boldsymbol{t,k}}^{(i)}) = \pi(\mathbf{y}_t^{(i)}|\mathbf{x}_{t,k}^{(i)}, \boldsymbol{\xi})$. Also, the bootstrap filter allows any error distribution as long as $\pi(\mathbf{y}_t^{(i)}|\mathbf{x}_{t,k}^{(i)}, \boldsymbol{\xi})$ can be evaluated pointwisely. However, propagating according to the transition density is often inefficient if the model is strongly stochastic such as in the Schlögl-model (Fig. 3B). This is because a large number of particles may need to be used to capture stochastic events. In these scenarios, making the proposal distribution $q(\cdot)$ depend on data is necessary for efficient simulation. In this sense, having "guided proposals" that at time $t-1$ propagate the particles for states $\mathbf{x}_{t-1}$ conditionally on the next data point $\mathbf{y}_t$ can be much more efficient [S19, S20, S21]. However, guided proposals are typically limited to specific error-models.

For propagators using the transition density (i.e. the bootstrap filter), in our framework the particles can be propagated using the SSA direct method [S7], the Extrande-method [S22], tau-leaping method (Eq. S5) and the Langevin-method (Eq. S7). The method to choose depends on which leap-conditions are fulfilled (Eq. S4 and S6).

Regarding guided proposals, our framework currently includes the modified diffusion for the Langevin-method [S23]. First assume a linear error model

$$\mathbf{y}_t^{(i)} = \mathbf{P}\mathbf{x}_t^{(i)} + \boldsymbol{\epsilon}, \quad \boldsymbol{\epsilon} \sim \mathcal{N}(0, \boldsymbol{\Sigma}), \tag{S15}$$

where $\mathbf{P}$ is a constant matrix (as an example, if $\mathbf{P}$ is diagonal with diagonal entries in $\{0, 1\}$, then a 0 on the diagonal would denote a corresponding unobserved coordinate of $\mathbf{x}$). Here also assume that the model dynamics are described by the Langevian-equation, and an appropriate time-discretisation $t_l = \tau_0 < \cdots < \tau_m = t_{l+1}$ where $\Delta\tau = \tau_{k+1} - \tau_k$. Then a linear Gaussian approximation of the conditional distribution $q(\mathbf{x}_{\tau_{k+1}}^{(i)}|\mathbf{y}_t^{(i)}, \mathbf{x}_{\tau_k}^{(i)}, \mathbf{c}^{(i)}, \boldsymbol{\kappa}, \boldsymbol{\xi})$, is given by

$$\begin{aligned}
&q(\mathbf{x}_{\tau_{k+1}}^{(i)}|\mathbf{y}_t^{(i)}, \mathbf{x}_{\tau_k}^{(i)}, \mathbf{c}^{(i)}, \boldsymbol{\kappa}, \boldsymbol{\xi}) \sim \mathcal{N}\Big(\mathbf{x}_{\tau_k}^{(i)} + \boldsymbol{\mu}(\mathbf{x}_{\tau_k}^{(i)}, \mathbf{c}^{(i)}\boldsymbol{\kappa})\Delta\tau, \boldsymbol{\Psi}(\mathbf{x}_{\tau_k}^{(i)}, \mathbf{c}^{(i)}\boldsymbol{\kappa})\Delta\tau\Big) \quad \text{where} \\
&\boldsymbol{\mu} = \boldsymbol{\alpha}_{\tau_k} + \boldsymbol{\beta}_{\tau_k}\mathbf{P}(\mathbf{P}^T\boldsymbol{\beta}_{\tau_k}\mathbf{P}\Delta_k + \Sigma)^{-1}(\mathbf{y}_t - \mathbf{P}^T(\mathbf{x}_{\tau_k} + \alpha_{\tau_k}\Delta_k) \quad \text{and} \\
&\boldsymbol{\Psi} = \boldsymbol{\beta}_{\tau_k} - \boldsymbol{\beta}_{\tau_k}\mathbf{P}(\mathbf{P}^T\boldsymbol{\beta}_{\tau_k}\mathbf{P}\Delta_k + \Sigma)^{-1}\mathbf{P}^T\boldsymbol{\beta}_{\tau_k}\Delta\tau,
\end{aligned} \tag{S16}$$

where $\boldsymbol{\alpha}_{\tau_k} = \boldsymbol{\alpha}(\mathbf{x}_{\tau_k}^{(i)}, \mathbf{c}^{(i)}\boldsymbol{\kappa})$, $\boldsymbol{\beta}_{\tau_k} = \boldsymbol{\beta}(\mathbf{x}_{\tau_k}^{(i)}, \mathbf{c}^{(i)}\boldsymbol{\kappa})$ and $\Delta_k = t - \tau_k$. Overall, this allows the particles to be propagated according to

$$\mathbf{x}_{\tau_{k+1}}^{(i)} \approx \mathbf{x}_{\tau_k}^{(i)} + \boldsymbol{\mu}(\mathbf{x}_{\tau_k}^{(i)}, \mathbf{c}^{(i)}\boldsymbol{\kappa})\Delta\tau + \sqrt{\boldsymbol{\Psi}(\mathbf{x}_{\tau_k}^{(i)}, \mathbf{c}^{(i)}\boldsymbol{\kappa})\Delta\tau}\mathbf{u}, \quad \mathbf{u} \sim \mathcal{N}(0, \mathbf{I}_{P\times P}). \tag{S17}$$

A guided proposal, such as the modified diffusion bridge, produces filters that are often statistically more efficient than transition-density based propagators, for the purpose of approximating integrals such as those defining likelihood terms [S23, S24]. However, the guided proposal outlined above is limited to the stated linear error model in Eq. S15 and to models having Gaussian transitions densities (for example, the Euler-Maruyama discretisation induces a Gaussian approximation to the true transition density).

Lastly, there are additional propagators to those mentioned here. The modular nature of our code makes it straightforward to implement additional ones, such as guided tau-leaping proposals [S25].

---

**Algorithm 2** Gibbs-sampler

---

**Input** Observed data $\mathbf{y}$, initial values and priors for $\boldsymbol{\kappa}, \boldsymbol{\eta}, \boldsymbol{\xi}$ and number of samples $n_{\text{samp}}$
**Output** Posterior distribution $\{\mathbf{c}, \boldsymbol{\kappa}, \boldsymbol{\eta}, \boldsymbol{\xi}\}_{j=1}^{n_{\text{samp}}}$

1: Initialisation: Set $\mathbf{c}^{(i,0)}$ to the mean of $\pi(\mathbf{c}|\boldsymbol{\eta}^{(0)})$. Draw $\mathbf{u}^{(i,0)} \sim g(\cdot)$ and estimate the initial observed likelihood value $\hat{\pi}_{\mathbf{u}^{(i)}}(\mathbf{y}^{(i)}|\mathbf{c}^{(i,0)}, \boldsymbol{\kappa}^{(0)}, \boldsymbol{xi}^{(0)})$ for $i = 1, \ldots, M$. Set iteration counter $j = 1$.

2: Update cell-individual parameters $\mathbf{c}^{(i)}$ for $i = 1, \ldots, M$:

a: Propose random numbers for estimating the likelihood $\mathbf{u}^{(i,*)} \sim g(\cdot)$

b: Propose new cell-individual parameters $\mathbf{c}^{(i,*)} \sim q(\cdot|\mathbf{c}^{(i,j-1)})$

c: Compute likelihood $\hat{\pi}_{\mathbf{u}^{(i,*)}}(\mathbf{y}^{(i)}|\mathbf{c}^{(i,*)}, \boldsymbol{\kappa}^{(j-1)}, \boldsymbol{\eta}^{(j-1)}, \boldsymbol{\xi}^{(j-1)})$ via Alg. 1.

d: Accept $\mathbf{u}^{(i,*)}$ and $\mathbf{c}^{(i,*)}$ with probability

$$\min\left\{1, \frac{\pi(\mathbf{c}^{(i,*)}|\boldsymbol{\eta}^{(j-1)})}{\pi(\mathbf{c}^{(i,j-1)}|\boldsymbol{\eta}^{(j-1)})} \cdot \frac{\hat{\pi}_{\mathbf{u}^{(i,*)}}(\mathbf{y}^{(i)}|\mathbf{c}^{(i,*)}, \boldsymbol{\kappa}^{(j-1)}, \boldsymbol{\xi}^{(j-1)})}{\hat{\pi}_{\mathbf{u}^{(i,j-1)}}(\mathbf{y}^{(i)}|\mathbf{c}^{(i,j-1)}, \boldsymbol{\kappa}^{(j-1)}, \boldsymbol{\xi}^{(j-1)})} \cdot \frac{q(\mathbf{c}^{(i,j-1)}|\mathbf{c}^{(i,*)})}{q(\mathbf{c}^{(i,*)}|\mathbf{c}^{(i,j-1)}})\right\}$$

and set $\mathbf{c}^{(i,j)} = \mathbf{c}^{(i,*)}$ and $\mathbf{u}^{(i,j)} = \mathbf{u}^{(i,*)}$ if accept, else $\mathbf{c}^{(i,j)} = \mathbf{c}^{(i,j-1)}$ and $\mathbf{u}^{(i,j)} = \mathbf{u}^{(i,j-1)}$.

3: Update cell-common parameters $\boldsymbol{\kappa}$ and $\boldsymbol{\xi}$

a: Propose new cell-common parameters $(\boldsymbol{\kappa}^{(*)}, \boldsymbol{xi}^{(*)}) \sim q(\cdot|\boldsymbol{\kappa}^{(j-1)}, \boldsymbol{xi}^{(j-1)})$.

b: Compute likelihood $\hat{\pi}_{\mathbf{u}^{(j)}}(\mathbf{y}|\mathbf{c}^{(j)}, \boldsymbol{\kappa}^{(*)}, \boldsymbol{\xi}^{(*)}) =$
$\prod_{i=1}^{M} \hat{\pi}_{\mathbf{u}^{(i,j)}}(\mathbf{y}^{(i)}|\mathbf{c}^{(i,j)}, \boldsymbol{\kappa}^{(*)}, \boldsymbol{\xi}^{(*)})$ by running Alg. 1 for individuals $i = 1, \ldots, M$.

c: Accept $(\boldsymbol{\kappa}^{(*)}, \boldsymbol{\xi}^{(*)})$ with probability

$$\min\left\{1, \frac{\pi(\boldsymbol{\kappa}^{(*)}, \boldsymbol{\xi}^{(*)})}{\pi(\boldsymbol{\kappa}^{(j-1)}, \boldsymbol{\xi}^{(j-1)})} \cdot \frac{\hat{\pi}_{\mathbf{u}^{(j)}}(\mathbf{y}|\mathbf{c}^{(j)}, \boldsymbol{\kappa}^{(*)}, \boldsymbol{\xi}^{(*)})}{\hat{\pi}_{\mathbf{u}^{(j)}}(\mathbf{y}|\mathbf{c}^{(j)}, \boldsymbol{\kappa}^{(j-1)}, \boldsymbol{\xi}^{(j-1)}} \cdot \frac{q(\boldsymbol{\kappa}^{(j-1)}, \boldsymbol{\xi}^{(j-1)}|\boldsymbol{\kappa}^{(*)}, \boldsymbol{\xi}^{(*)})}{q(\boldsymbol{\kappa}^{(*)}, \boldsymbol{\xi}^{(*)}|\boldsymbol{\kappa}^{(j-1)}, \boldsymbol{\xi}^{(j-1)})}\right\}$$

and set $(\boldsymbol{\kappa}^{(j)}, \boldsymbol{\xi}^{(j)}) = (\boldsymbol{\kappa}^{(*)}, \boldsymbol{\xi}^{(*)})$ if accept, else $(\boldsymbol{\kappa}^{(j)}, \boldsymbol{\xi}^{(j)}) = (\boldsymbol{\kappa}^{(j-1)}, \boldsymbol{\xi}^{(j-1)})$.

4: Update population parameters $\boldsymbol{\eta}$ by running an HMC-sampler that targets

$$\pi(\boldsymbol{\eta}|\mathbf{c}^{(j)}, \boldsymbol{\kappa}^{(j)}, \boldsymbol{\xi}^{(j)}, \mathbf{y}) \propto \pi(\boldsymbol{\eta}) \prod_{i=1}^{M} \pi(\mathbf{c}^{(i,j)}|\boldsymbol{\eta})$$

5: If $j = n_{\text{samp}}$ stop. Else, set $j = j + 1$ and go to step 2.

---

#### S2.3 Gibbs-sampler

We consider two Gibbs samplers: the first one is described in this section, while the second (and particularly efficient) one is in section S2.4. Our Gibbs sampler targets the posterior in Eq. S2 by sampling from the conditionals (see the "blocked Gibbs" in [S2])

$$
\begin{aligned}
&1.\ \pi(\mathbf{c}^{(i)}, \mathbf{u}^{(i)} | \boldsymbol{\kappa}, \boldsymbol{\eta}, \boldsymbol{\xi}, \mathbf{y}) \propto \pi(\mathbf{c}^{(i)} | \boldsymbol{\eta}) \hat{\pi}_{\mathbf{u}^{(i)}}(\mathbf{y}^{i} | \mathbf{c}^{(i)}, \boldsymbol{\kappa}, \boldsymbol{\xi}) g(\mathbf{u}^{(i)}), \qquad i = 1, \dots, M \\
&2.\ \pi(\boldsymbol{\kappa}, \boldsymbol{\xi} | \mathbf{c}, \boldsymbol{\eta}, \mathbf{y}, \mathbf{u}) \propto \pi(\boldsymbol{\kappa}, \boldsymbol{\xi}) \prod_{i=1}^{M} \hat{\pi}_{\mathbf{u}^{(i)}}(\mathbf{y}^{(i)} | \mathbf{c}^{(i)}, \boldsymbol{\kappa}, \boldsymbol{\xi}) \\
&3.\ \pi(\boldsymbol{\eta} | \mathbf{c}, \boldsymbol{\kappa}, \boldsymbol{\xi}, \mathbf{y}, \mathbf{u}) \propto \pi(\boldsymbol{\eta}) \prod_{i=1}^{M} \pi(\mathbf{c}^{(i)} | \boldsymbol{\eta}).
\end{aligned} \tag{S18}
$$

The pseudo-code is presented in Algorithm 2. Briefly, the individual parameters $\mathbf{c}^{(i)}$ and the cell-common parameters $(\boldsymbol{\kappa}, \boldsymbol{\xi})$ are updated via a pseudo-marginal approach since their conditionals depends on the observed data likelihood (notice, in step 1 we perform $M$ samplings, one for each $\mathbf{c}^{(i)}$). Here, $\mathbf{u}^{(i)}$ are the auxiliary-variables used to obtain an unbiased estimate of the likelihood via a particle filter (Alg. 1). For the Langevin and Poisson-methods we can induce a high correlation in these auxiliary variables to reduce the number of particles used. Lastly, to propose parameter efficiently, adaptive kernels $q(\cdot)$ are employed for each individual [S26, S27, S28]. Notice, new pseudorandom variates $\boldsymbol{u}$ are created in step 2a of Alg. 2, however the $\boldsymbol{u}$ that are used in step 3b are the ones returned from step 2d, that is we recycle the pseudorandom variates in step 3b when evaluating the approximate likelihood $\hat{\pi}_{\mathbf{u}^{(j)}}(\mathbf{y} | \mathbf{c}^{(j)}, \boldsymbol{\kappa}^{(*)}, \boldsymbol{\xi}^{(*)})$ at the newly proposed $(\boldsymbol{\kappa}^{(*)}, \boldsymbol{\xi}^{(*)})$.

The population parameters $\boldsymbol{\eta}$ are proposed using an efficient Hamiltonian Monte Carlo (HMC) sampler [S3, S4, S5]. This allows for a wide range of different prior distributions; $\mathbf{c}^{(i)} \sim \pi(\boldsymbol{\eta})$. Currently, a log-normal distribution is implemented: $\mathbf{c}^{(i)} \sim \mathcal{LN}(\boldsymbol{\mu}, \boldsymbol{\Omega})$ where $\boldsymbol{\mu}$ and $\boldsymbol{\Omega}$ are the mean and covariance matrix of the associate Gaussian distribution. Here $\boldsymbol{\Omega}$ can be diagonal, or non-diagonal of the form $\boldsymbol{\Omega} = \mathrm{D}(\boldsymbol{\tau})\boldsymbol{\Phi}\mathrm{D}(\boldsymbol{\tau})$ where $\mathrm{D}(\boldsymbol{\tau})$ is a diagonal matrix with the scale-vector $\boldsymbol{\tau}$ on the diagonal, and $\boldsymbol{\Phi}$ is a non-diagonal correlation matrix. Typically, this separation approach performs well for covariance matrix inference [S29]. Additionally, the modular structure of the code allows the user to add additional distributions for $\mathbf{c}^{(i)}$. Moreover, for models where the HMC-sampler might be to computationally demanding (non-diagonal $\boldsymbol{\Omega}$ with many individuals) a conjugate-prior update scheme can be implemented [S29].

#### S2.4 Modified Gibbs-sampler

The Gibbs-sampler (Alg. 2) produces exact Bayesian inference, however, the run-time can be substantial. A contributing factor is that sampling of $(\boldsymbol{\kappa}, \boldsymbol{\xi})$ requires a stochastic approximation of the product of the individual likelihoods (Alg. 2 step 3). As we approximate this product by computing a stochastic approximation of the likelihood for each individual, the variance of the approximation is typically large. As discussed in section S2.5 a “large” variance may considerably reduce the chain mixing and even making the chain stuck. To avoid this, a possibility is to use a larger number of particles for each individual ($N^{(i)}$ Alg. 1), which would however result in longer run-times.

To reduce the number of particles, and thus the run-time, we introduced a “modified sampler” (Alg. 3). Here, inspired by how cell-constant parameters can be handled in the software Monolix [S30], we perturb the original SSMEM slightly by treating the cell-common parameters $(\boldsymbol{\kappa}, \boldsymbol{\xi})$ as parameters that vary between cells. However, we fix their variance at a small value. For example, we may assume that $(\boldsymbol{\kappa}^{(i)}, \boldsymbol{\xi}^{(i)}) \sim \mathcal{N}\big((\boldsymbol{\kappa}_{pop}, \boldsymbol{\xi}_{pop}), \delta \cdot \mathbf{I}\big)$. Here, $\delta > 0$ is a small tuning parameter specified by the user. We set $\delta$ somewhat arbitrarily, but we found that for parameters having magnitude $10^0 - 10^1$ a value of $\delta = 0.01$ worked well (Fig. S5). In scenarios where the magnitude of $(\boldsymbol{\kappa}, \boldsymbol{\xi})$ is unknown, a pilot-run can be performed to find the scale of these parameters, and $\delta$ adjusted accordingly (pilot runs need typically to be performed in any case, e.g. when observing if adaptive MCMC works adequately well). Ultimately, a "modified Gibbs sampler" is a standard Gibbs sampler, but performed on a perturbed SSMEM, where the perturbation is induced by the use of $\delta$. Overall, given this perturbation of the SSMEM, the focus for the inference becomes to infer $(\boldsymbol{\xi}_{pop}, \boldsymbol{\kappa}_{pop})$ alongside with the population parameters $\boldsymbol{\eta}$. Overall, as we update $(\boldsymbol{\kappa}_{pop}, \boldsymbol{\xi}_{pop})$ using an HMC-sampler the product of the individual likelihoods is not required for any parameter update. Rather, only the individual likelihoods are used in the pseudo-marginal approach. Thus, only the variance of the individual likelihoods needs to be controlled, and this requires fewer particles compared to controlling the variance of the population likelihood (i.e. the product of the individual likelihoods). Moreover, as described in S2.5, this allows us to tune the number of particles per individual. In summary, the perturbation induced by $\delta$ allows $(\boldsymbol{\xi}_{pop}, \boldsymbol{\kappa}_{pop})$ to be inferred alongside with the population parameters $\boldsymbol{\eta}$ in step 3 of the Gibbs sampler, and $(\boldsymbol{\xi}^{(i)}, \boldsymbol{\kappa}^{(i)})$ to be inferred alongside $\mathbf{c}^{(i)}$ in step 1. Step 2 is thus avoided.

As noted for the Schlögl model (Fig. S3) the modified sampler is substantially faster. However, as we can see for the Ornstein-Uhlenbeck model this is at the price of $(\boldsymbol{\kappa}_{pop}, \boldsymbol{\xi}_{pop})$ having slightly wider credibility intervals (Fig. S5). However, we consider this a worthwhile compromise. Moreover, if the user is interested in exact inference, the modified sampler can be used for pilot-runs. Then the standard sampler (Alg. 2) can be run from the posterior-mode obtained from the modified sampler. Lastly, as no parameters are likely to be constant between individuals [S31], the modified sampler in a sense infers a more realistic model. However, how more realistic is hard to quantify since $\delta$ still is determined quite arbitrarily.

#### S2.5 The number of particles in the Gibbs-samplers

Choosing an appropriate number of particles for the pseudo-marginal steps of the Gibbs-samplers is crucial for performance. If too few particles are used the Gibbs-sampler is likely to get stuck. This is because the estimated likelihood $\hat{Z}$ will have a large variance. More precisely, imagine a parameter $\boldsymbol{\theta}_1$ is accepted with an unusually high likelihood value (and the latter is also virtually stored, since it is a function of the accepted $\boldsymbol{u}$ and accepted parameters) owing to the variability of the estimated likelihood. Now, it is likely that for a newly proposed $\boldsymbol{\theta}^*$ the corresponding estimated likelihood will have a smaller value than the previously accepted likelihood. Overall, a long series of rejections may be started this way. Moreover, the parameters region targeted by adaptive MCMC proposals may result as drastically being shrunken, as many adaptive schemes reduce the size of their proposal region following a rejection [S27, S28]. All-in-all, although theoretically any number of particles may be selected for the Gibbs sampler to return exact Bayesian inference (in view of the properties of pseudo-marginal methods) as the number of iterations go to infinity, in practice a "large enough" amount of particles is crucial [S32, S15, S33, S34]. However, the run-time of the particle filter increases with the number of particles. Therefore we need to find a trade-off. Here, principled approaches for setting the number of particles are covered.

##### S2.5.1 Correlating the particles

The number of particles in the Gibbs-samplers can be reduced by employing correlated particle filters. In the Gibbs-samplers an unbiased likelihood estimate is obtained by proposing auxiliary variables $\mathbf{u}^{(i)}$ from an independent proposal kernel $g(\cdot)$ (e.g a Normal-distribution). However, as discussed in [S35] the proposal kernel can also be of the form $K(\mathbf{u}^{i*}|\mathbf{u}^{(i)})$, with

**Algoritm 3** Modified Gibbs-sampler

**Input** Observed data $\mathbf{y}$, initial values and priors for $\boldsymbol{\kappa}, \boldsymbol{\eta}, \boldsymbol{\xi}$, perturbing variance $\delta$ and number of samples $n_{\text{samp}}$

**Output** Posterior distribution $\{\mathbf{c}, \boldsymbol{\kappa}_{pop}, \boldsymbol{\eta}, \boldsymbol{\xi}_{pop}\}_{j=1}^{n_{\text{samp}}}$

1: Initialise: Put $\mathbf{c}^{(i,0)}$ to the mean of $\pi(\mathbf{c}|\boldsymbol{\eta}^{(0)})$. Draw $\mathbf{u}^{(i,0)} \sim g(\cdot)$ and estimate initial observed likelihood value $\hat{\pi}_{\mathbf{u}^{(i,0)}}(\mathbf{y}^{(i)}|\mathbf{c}^{(i,0)}, \boldsymbol{\kappa}^{(0)}, \boldsymbol{\xi}^{(0)})$ for $i = 1, \dots, M$. Set iteration counter $j = 1$.

2: Update cell-individual parameters $\mathbf{c}^{(i)}$ for $i = 1, \dots, M$:

a: Propose new auxiliary variables for estimating likelihood $u^{(i,*)} \sim g(\cdot)$.

b: Propose new cell-individual parameters $\mathbf{c}^{(i,*)} \sim q(\cdot|\mathbf{c}^{(i,j-1)})$.

c: Propose new cell-individual $(\boldsymbol{\kappa}^{(i,*)}, \boldsymbol{\xi}^{(i,*)}) \sim q(\cdot|\boldsymbol{\kappa}^{(i,j-1)}, \boldsymbol{\xi}^{(i,j-1)})$.

d: Compute likelihood $\hat{\pi}_{\mathbf{u}^{(i,*)}}(\mathbf{y}^{(i)}|\mathbf{c}^{(i,*)}, \boldsymbol{\kappa}^{(i,*)}, \boldsymbol{\xi}^{(i,*)})$ via Alg. 1.

e: Accept $\mathbf{u}^{(i,*)}$, $\mathbf{c}^{(i,*)}$, and $(\boldsymbol{\kappa}^{(i,*)}, \boldsymbol{\xi}^{(i,*)})$ with probability

$$\min\Bigg\{1, \frac{\pi(\mathbf{c}^{(i,*)}|\boldsymbol{\eta}^{(j-1)})}{\pi(\mathbf{c}^{(i,j-1)}|\boldsymbol{\eta}^{(j-1)})} \cdot \frac{\hat{\pi}_{\mathbf{u}^{(i,*)}}(\mathbf{y}^{(i)}|\mathbf{c}^{(i,*)}, \boldsymbol{\kappa}^{(i,*)}, \boldsymbol{\xi}^{(i,*)})}{\hat{\pi}_{\mathbf{u}^{(i,j-1)}}(\mathbf{y}^{(i)}|\mathbf{c}^{(i,j-1)}, \boldsymbol{\kappa}^{(i,j-1)}, \boldsymbol{\xi}^{(i,j-1)})} \cdot \frac{q(\mathbf{c}^{(i,j-1)}|\mathbf{c}^{(i,*)})}{q(\mathbf{c}^{(i,*)}|\mathbf{c}^{(i,j-1)})} \cdot \frac{\pi(\boldsymbol{\kappa}^{(i,*)}, \boldsymbol{\xi}^{(i,*)}|\boldsymbol{\kappa}_{pop}^{(j-1)}, \boldsymbol{\xi}_{pop}^{(j-1)})}{\pi(\boldsymbol{\kappa}^{(i,j-1)}, \boldsymbol{\xi}^{(i,j-1)}|\boldsymbol{\kappa}_{pop}^{(j-1)}, \boldsymbol{\xi}_{pop}^{(j-1)})} \cdot \frac{q(\boldsymbol{\kappa}^{(i,j-1)}, \boldsymbol{\xi}^{(i,j-1)}|\boldsymbol{\kappa}^{(i,*)}, \boldsymbol{\xi}^{(i,*)})}{q(\boldsymbol{\kappa}^{(i,*)}, \boldsymbol{\xi}^{(i,*)}|\boldsymbol{\kappa}^{(i,j-1)}, \boldsymbol{\xi}^{(i,j-1)})}\Bigg\}$$

and set $\mathbf{c}^{(i,j)} = \mathbf{c}^{(i,*)}$, $(\boldsymbol{\kappa}^{(i,j)}, \boldsymbol{\xi}^{(i,j)}) = (\boldsymbol{\kappa}^{(i,*)}, \boldsymbol{\xi}^{(i,*)})$ and $\mathbf{u}^{(i,j)} = \mathbf{u}^{(i,*)}$ if accept, else $\mathbf{c}^{(i,j)} = \mathbf{c}^{(i,j-1)}$, $(\boldsymbol{\kappa}^{(i,j)}, \boldsymbol{\xi}^{(i,j)}) = (\boldsymbol{\kappa}^{(i,j-1)}, \boldsymbol{\xi}^{(i,j-1)})$ and $\mathbf{u}^{(i,j)} = \mathbf{u}^{(i,j-1)}$.

3: Update population parameters $\boldsymbol{\eta}$ by running an HMC-sampler that targets

$$\pi(\boldsymbol{\eta}|\mathbf{c}^{(j)}, \boldsymbol{\kappa}^{(j)}, \boldsymbol{\xi}^{(j)}, \boldsymbol{\kappa}_{\text{pop}}^{(j)}, \boldsymbol{\xi}_{\text{pop}}^{(j)}, \mathbf{y}, \mathbf{u}^{(j)}) \propto \pi(\boldsymbol{\eta}) \prod_{i=1}^{M} \pi(\mathbf{c}^{(i,j)}|\boldsymbol{\eta}).$$

4: Update cell-common parameters $(\boldsymbol{\kappa}_{pop}, \boldsymbol{\xi}_{pop})$ by running a HMC-sampler that targets

$$\pi(\boldsymbol{\kappa}_{\text{pop}}, \boldsymbol{\xi}_{\text{pop}}|\mathbf{c}^{(j)}, \boldsymbol{\kappa}^{(j)}, \boldsymbol{\xi}^{(j)}, \boldsymbol{\eta}^{(j)}, \mathbf{y}, \mathbf{u}^{(j)}) \propto \pi(\boldsymbol{\kappa}_{\text{pop}}, \boldsymbol{\xi}_{\text{pop}}) \prod_{i=1}^{M} \pi(\boldsymbol{\kappa}^{(i,j)}, \boldsymbol{\xi}^{(i,j)}|\boldsymbol{\kappa}_{pop}, \boldsymbol{\xi}_{pop})$$

where

$$\pi(\boldsymbol{\kappa}^{(i,j)}, \boldsymbol{\xi}^{(i,j)}|\boldsymbol{\kappa}_{pop}, \boldsymbol{\xi}_{pop}) = \mathcal{N}\big((\boldsymbol{\kappa}_{pop}, \boldsymbol{\xi}_{pop}), \delta\mathbf{I}\big), \quad \delta > 0.$$

5: If $j = n_{\text{samp}}$ stop. Else, set $j = j + 1$ and go to step 2.

$$K(\mathbf{u}^{(i,*)}|\mathbf{u}^{(i)}) = \mathcal{N}\big(\mathbf{u}^{(i,*)}; \rho\mathbf{u}^{(i)}, (1-\rho^2)\mathbf{I}_d\big), \tag{S19}$$

where $\mathbf{I}_d$ is the identity matrix and $\rho \in [0.95, 0.999]$ which induces a strong positive correlation between successive approximations to the likelihood, $\hat{Z}^*$ and $\hat{Z}$. By inducing a strong-correlation the variance of the acceptance ratio (step 2d and step 3c in Alg. 2) is reduced. While this does not reduce the variance of the approximated likelihoods, the intuition is that what matters is reducing the variability of the likelihoods *ratio*. Consequently, fewer particles can be used without the risk of the sampler getting stuck.

Three factors must be accounted for when employing correlated particles. Firstly, to correlate the particles the amount of auxiliary variables $\mathbf{u}^{(i)}$ must be known a-priori. For the tau-leaping-method and Langevian-method the number of random-numbers (auxiliary variables) used for running the particle filter is known given a simulation discretisation stepsize $\tau$, however, for the Gillespie's method the number of auxiliary variables is itself a random variable, and thus a correlated particle filter cannot be employed. Secondly, the auxiliary variables generated by Eq. S19 are normally distributed. However, the resampling steps in the particle filter require uniform random numbers, and the tau-leaping-method requires Poisson-random numbers. In these cases uniform-numbers can be obtained via the CDF of the normal distribution $\Phi(\mathbf{u}^{(i)})$. Moreover, for a given uniform draw we have implemented an efficient Poisson-generator (`https://github.com/cvijoviclab/PEPSDI/blob/main/Code/Stochastic_solvers/Poisson.jl`). Thirdly, the re-sampling step can break the correlation between particles. This can be alleviated by using an Euclidean sorting step [S36]. However, this sorting procedure can fail for larger measurements errors as a high noise-level can cause correlations to deteriorate [S36].

##### S2.5.2 Tuning the particles

In the Gibbs-samplers the number of particles to use can be tuned. Here, it is important to perform the tuning at a central-posterior location for the parameters to infer. In fact, compared to when computed at a parameter close to a mode, the estimated likelihood has a larger variance than when computed far away from a mode. This is because conditionally to implausible parameter values the particles will often end-up far away from the data, thus providing a poor approximation to the likelihood (this is informally discussed in `https://darrenjw.wordpress.com/2014/06/08/tuning-particle-mcmc-algorithms/`). Since the goal is to perform sampling around a central-posterior location, it is thus desirable to tune at a central-posterior location. To find such a location a pilot-run is required.

Overall, we tune the particles by first running a pilot run (with a high number of particles) for $3000 - 5000$ iterations to prevent the sampler from getting stuck. Then at the mode (central posterior location) we fix $\mathbf{c}^{(i)}, \boldsymbol{\kappa}, \boldsymbol{\xi}$ and choose the number of particles according to the correlation level $\rho$. When the particles are not correlated we choose $N^{(i)}$ such that the variance of the log-likelihood ($\sigma^2_{N^{(i)}}$) is smaller than 2. When the particles are correlated ($\rho \neq 0$) we follow [S2] and choose $N^{(i)}$ such that $\sigma^2_{N^{(i)}} < 2.16^2/(1-\rho_l^2)$, where $\rho_l$ is the estimated correlation between $\pi(\mathbf{y}^i, \mathbf{u}^i|\boldsymbol{\kappa}, \boldsymbol{\xi}, \mathbf{c}^{(i)})$ and $\pi(\mathbf{y}^{(i)}, \mathbf{u}^{(i,*)}|\boldsymbol{\kappa}, \boldsymbol{\xi}, \mathbf{c}^{(i)})$. For the standard Gibbs-sampler we use the same $N^{(i)}$ for all individual and we tune according to the full data likelihood (Alg. 2 step 3c). As we sidestep the full data likelihood for the modified sampler (default option in PEPSDI) we use the same number of particles for each individual in the pilot run, but, we then tune the particles based on the variance of the individuals likelihoods. This allows the particles to vary between individuals for the main inference run, and typically this results in fewer particles being required per individual. Naturally, for the modified sampler the particles can also, if necessary for computational efficiency, vary between individuals for the pilot run.

### S3 Guidelines for running our inference framework

Here we provide guidelines for how to run our inference framework for a user provided state-space model. Guidelines refer to both the default perturbed sampler (Alg. 3 where $\boldsymbol{\kappa}, \boldsymbol{\xi}$ are perturbed to weakly vary between cells) and the non-perturbed sampler (Alg. 2 where $\boldsymbol{\kappa}, \boldsymbol{\xi}$ are cell-constant). When appropriate, remarks are provided for the different steps.

1. Choose appropriate priors for $\boldsymbol{\eta}$, $\boldsymbol{\kappa}$ and $\boldsymbol{\xi}$.
   - Any distribution in the Julia distributions-package can be used.
2. If possible choose suitable starting values for $\boldsymbol{\eta}$, $\boldsymbol{\kappa}$ and $\boldsymbol{\xi}$.
   - The mean of the priors can be used if no suitable starting points is known.
3. Execute a pilot run with $3,000 - 5,000$ iterations using a large particle number for each individual (say $> 1,000$ particles).
   - A large particle number is necessary to prevent the sampler from getting stuck during the pilot run.
   - If the aim is to do inference using Gillespie's method, an approximate simulation algorithm can be used for the pilot run.
4. Tune the particles at the end location of the pilot run according to the tuning criteria.
   - Our implementation automatically tunes the particles at the end of the pilot-run.
   - Run time strongly depends on the number of particles, making this a crucial step.
   - The default perturbed sampler does not need to compute the full data likelihood (product of individual likelihoods). Consequently it tunes the particles on an individual basis following the pilot run (all individuals have same number of particles in the pilot). The non-perturbed sampler uses the same number of particles for each individual, and the particles are tuned to control the variance of the full data-likelihood often resulting in a large number of particles.
5. Run a further main inference by initializing the Gibbs sampler at the posterior mean obtained from the pilot run, and using the number of particles as obtained from step 4. Further, to propose $\mathbf{c}, \boldsymbol{\kappa}$ and $\boldsymbol{\xi}$ use the tuned covariance matrices obtained in the pilot run, as returned by an adaptive-MCMC scheme (Fig. 4).
   - Our implementation automatically loads the tuned covariance-matrices, posterior mean-values and number of particles.
6. Judge the quality of inference via diagnostic plots such as trace-plots and posterior visual checks.
   - If trace-plots shows that the sampler got stuck, increase the number of particles and re-run step 5. The tuning criteria, albeit typically working well, can fail.

**Remarks for step 2 and 5**: Firstly, the tau-leaping and Langevin simulators require a step-length to be set. This can be done by simulating the model at the starting values using different step-lengths, and then choose the maximum value from which the results do not appear to change. Secondly, use correlated particles and guided proposals to reduce the number of particles required. Thirdly, our implementation employs multi-threading to update the individual parameters in parallel. The user can set the number of threads via the Julia threads variable.

### S4 A tutorial on constructing a single-cell dynamic model

Single-cell dynamic models can elucidate cellular reaction dynamics and sources of cell-to-cell variability. To help both modellers and experimentalists utilise such models, we provide a brief tutorial on model construction. For the gene regulatory network in Fig. 2a, we describe how the reaction dynamics are formulated, how intrinsic and extrinsic noise can be modelled, and in a complementary notebook[S1], how PEPSDI infers unknown model quantities.

As a starting point, we assume that the single-cell protein time-lapse data for the gene regulatory network in Fig. 2b has been collected. From here, the first modelling step is to formulate a literature-informed hypothesis of the reaction dynamics, and subsequently draw a reaction map (Fig. 2a). This reaction map (model) should be sufficiently complex to capture core features, for example, when modelling a gene regulatory network this can correspond to excluding the finer details of transcription and translation (as in Fig. 2a).

Given a reaction map, the next step is to formulate the propensities for each cellar process (reaction) in the reaction map (often via laws of mass-action):

$$\mathbf{h} = \begin{bmatrix} h_1 & h_2 & h_3 & h_4 \end{bmatrix} = \left[ \underbrace{c_1\big(1+\sin(\omega t)\big)}_{\text{Transcription}}, \quad \underbrace{c_2\text{mRNA}}_{\text{mRNA degradation}}, \underbrace{c_3\text{mRNA}}_{\text{Translation}}, \quad \underbrace{c_4\text{Protein}}_{\text{Protein degradation}} \right]. \tag{S20}$$

Formally, the propensity $h_h$ measures the probability of reaction $j$ to occur and is thus instrumental for model simulations. The rate constants $\mathbf{c}^{(i)}$ are measures of reaction-strength, and typically describe several cellular-components. For example, the probability of translation to occur is proportional to the amount of mRNA and ribosomes: $h_3 \propto [mRNA][Ribosome]$. Assuming constant amount of ribosomes in a cell, given amount enters the rate constant $c_3 \propto [\textit{Ribosome}]$ to yield the propensity in Eq. S20. Furthermore, the rate-constants can be time ($t$) dependent. Consider the transcription reaction, modelled to be regulated by a circadian clock transcription factor (TF); $\tilde{c}_1 \propto [TF](t) \propto c_1\big(1+\sin(\omega t)\big)$, where $\omega = 2\pi/24$ emulates a circadian-clock cycle of 24 hours.

Given propensities, the model can be simulated via a stochastic simulator to capture intrinsic noise. For small number of molecule (e.g 2-10 molecules) exact simulators are accurate [S7, S8], while faster approximate simulator are often accurate for larger molecule numbers [S6, S9]. If the number of molecules are unknown, an approximate simulator can be used initially, and if some model components have few molecules one can switch to an exact simulator.

Extrinsic noise can be modelled by letting the initial values (mRNA, Protein), and/or rate-constants vary between cells. For motivating the latter, consider the translation propensity; $h_3 = c_3 mRNA \propto [mRNA][\textit{Ribosome}]$. Assuming that ribosome abundance varies between cells due to extrinsic noise, $c_3$ should also vary. Similarly, $(c_1, c_2, c_4)$ might vary between cells, but, not arbitrarily. Typically, the rate-parameters can be constrained by assuming that these follow probability distributions with an appropriate support. A log-normal distribution [S37, S38] is the most common assumption. In the example here, this would imply that rate constants $\mathbf{c}^{(i)}$ follow a log-normal distribution $(c_1^{(i)}, \ldots, c_4^{(i)}) \sim \mathcal{LN}(\boldsymbol{\mu}, \boldsymbol{\Omega})$, where $\boldsymbol{\mu}$ and $\boldsymbol{\Omega}$ are unknown population parameters that describe the distribution of $\mathbf{c}^{(i)} = (c_1^{(i)}, \ldots, c_4^{(i)})$.

In summary, to construct a single-cell model i) draw a reaction network, ii) for each reaction formulate the propensity, iii) capture intrinsic noise via exact stochastic simulators [S7, S8], for

[S1] `https://github.com/cvijoviclab/PEPSDI/blob/main/Code/Examples/Multiple_individual/Tutorial_PEPSDI.ipynb` small molecule numbers, or approximate simulator for larger molecule numbers [S9, S6], and iv) capture extrinsic noise by letting rate-constants and/or initial values vary between cells by following a probability distribution. However, model construction is only the first step. For example, typically we also want to investigate if a model can accurately describe the observed data, characterise its behaviour, and use it to predict new behaviour. To this end, we need to know the rate-constants $\mathbf{c}^{(i)}$, and how they vary between cells. Thus, to fully utilise a derived model, we can use PEPSDI to infer in a Bayesian fashion the rate-parameters ($\mathbf{c}^{(i)}$ for individual $i$), the strength of the measurement error ($\boldsymbol{\xi}$), and the population parameters ($\boldsymbol{\mu}, \boldsymbol{\Omega}$). Furthermore, albeit the rate-parameters are unknown we can sometimes have a prior idea of, for example, their magnitude. Since PEPSDI follows a Bayesian-framework, the user can incorporate prior information into the inference via parameter priors. In a complementary notebook, we show a code for the model we derived here and how to use PEPSDI to perform the inference.

### S5 Simulation examples

Here, the models used in the simulation examples (circadian clock regulated gene network, and Schlögl-model) are presented. For each model, the model equations, simulation options, and inference details are presented.

#### S5.1 Circadian clock regulated gene-network model

The circadian-clock model regulated gene network (Fig. 2) consists of four reactions

$$\begin{aligned} &\mathcal{R}_1 \quad \phi \xrightarrow{c_1\left(1+\sin(\omega t)\right)} \text{mRNA} \\ &\mathcal{R}_2 \quad \text{mRNA} \xrightarrow{c_2} \phi \\ &\mathcal{R}_3 \quad \text{mRNA} \xrightarrow{c_3} \text{Protein} \\ &\mathcal{R}_4 \quad \text{Protein} \xrightarrow{c_4} \phi \end{aligned} \tag{S21}$$

with $\omega = 2\pi/24$ to emulate a circadian-clock cycle of 24 hours. The associated stoichiometry matrix is

$$\mathbf{S} = \begin{bmatrix} -1 & 0 \\ 0 & -1 \\ 1 & 0 \\ 0 & 1 \end{bmatrix}, \tag{S22}$$

and the propensity (hazard) vectors is

$$\mathbf{h}(X_t, \mathbf{c}) = \left[c_1\left(1+\sin(\omega t)\right), \quad c_2\text{mRNA}, \quad c_3\text{mRNA}, \quad c_4\text{Protein}\right]^{\mathrm{T}}. \tag{S23}$$

##### S5.1.1 Simulation of data-set

Using the Extrande-method [S8] 40 individuals were simulated using the circadian clock-model. For each individual, we simulated data at 64 equidistant time-points from 0.5 to 48 hours. Regarding the observations $Y_t$, we assumed that only the protein was observed. We also assumed that the observations were noise-corrupted by a normal noise; $Y_t = \text{Protein}_t + \epsilon_t$ where $\epsilon_t \sim \mathcal{N}(0, \xi^2) = \mathcal{N}(0, 2^2)$. The individual parameters $\mathbf{c} = (c_1, \ldots, c_4)$ were generated by applying the exponential function on random draws from a multivariate normal $\mathcal{N}(\boldsymbol{\mu}, \boldsymbol{\Omega})$, where $\boldsymbol{\Omega} = \mathrm{D}(\boldsymbol{\tau})\boldsymbol{\Phi}\mathrm{D}(\boldsymbol{\tau})$, where $\mathrm{D}(\cdot)$ refers to diagonal matrix with $\boldsymbol{\tau}$ on the diagonal. Here $\boldsymbol{\tau}$ is the scale-vector (variance), and $\boldsymbol{\Phi}$ is a non-diagonal correlation matrix. As the exponential of a normal-random variable is log-normal random variable, $\mathbf{c}$ becomes multivariate log-normal. The actual population parameters used where

$$\boldsymbol{\mu} = \begin{bmatrix} \log 3 \\ \log 30 \\ \log 3 \\ \log 2 \end{bmatrix} \approx \begin{bmatrix} 1.09 \\ 3.40 \\ 1.09 \\ 0.69 \end{bmatrix}, \quad \boldsymbol{\tau} = \begin{bmatrix} 0.4 \\ 0.2 \\ 0.2 \\ 0.1 \end{bmatrix},$$
$$\boldsymbol{\Phi} = \begin{bmatrix} 1.0 & -0.159014 & -0.175134 & -0.175134 \\ \cdot & 1.0 & -0.261888 & -0.22718 \\ \cdot & \cdot & 1.0 & -0.49255 \\ \cdot & \cdot & \cdot & 1.0 \end{bmatrix}. \quad \text{(S24)}$$

Note that $\boldsymbol{\Phi}$ is a symmetric positive-definite matrix.

#### S5.2 Inference details

The parameters inferred were the individual parameters $\mathbf{c}^{(i)}$, the mean population vector $\boldsymbol{\mu}$, the scale-vector $\boldsymbol{\tau}$ of the covariance matrix, the correlation matrix $\boldsymbol{\Phi}$, and the strength of the measurement error parameter $\xi$. Note, the covariance matrix was parameterised using the separation approach as $\mathrm{D}(\boldsymbol{\tau})\boldsymbol{\Phi}\mathrm{D}(\boldsymbol{\tau})$. This approach typically performs better than using conjugate priors [S29]. Furthermore, the individual parameters were inferred on the log-scale as they are positive.

A $\mathcal{N}(0, 2^2)$ prior was used for $\mu_1$. For $\mu_2, \mu_3, \mu_4$ we used weakly informative priors, given by $\mathcal{N}(0, 5^2)$, $\mathcal{N}(0, 5^2)$, $\mathcal{N}(0, 5^2)$ respectively. Note that none of this priors are centred around the true-values. Following the Stan user guide [S39], a half-Cauchy prior was applied on the scale-parameters, $\tau \sim \mathcal{C}(0, 2.5)$ where $\tau > 0$. A LKJ-prior with shape-parameter 4 was used for the correlation matrix $\boldsymbol{\Phi}$. Lastly, a gamma$(1, 2)$ prior was used for the error parameter $\xi$.

We ran the inference using the default options in PEPSDI, where the cell-constant parameters are slightly perturbed. We used as starting values a random sampled vector from the prior. The inference was run according to the guidelines in S3. Briefly, for the pilot run $5,000$ iterations were performed using 2,000 particles. The high number of particles were employed to control the variance of the particle filter estimated likelihood. At the end location of the pilot run, according to the criteria for non-correlated particle filters, we tuned the number of particles per individual, and the minimum, median and maximum number of particles required for an individual was 90, 245 and 1150 respectively. Notice, that we could not correlate the particles since when performing inference using the Extrande-simulator we do not, a priori, know the number of random numbers required by the particle filter. Lastly, starting from where the pilot-run ended, $50,000$ further iterations of the perturbed Gibbs sampler were run to produce the final inference.

#### S5.3 The Schlögl-model

The Schlögl-model [S40] consists of four reactions

$$\begin{aligned} &\mathcal{R}_1 \quad A + 2X \xrightarrow{\tilde{c}_1} 3X \\ &\mathcal{R}_2 \quad 3X \xrightarrow{c_2} A + 2X \\ &\mathcal{R}_3 \quad B \xrightarrow{\tilde{c}_3} X \\ &\mathcal{R}_4 \quad X \xrightarrow{c_4} B, \end{aligned} \quad \text{(S25)}$$

notice $\tilde{c}_1$ and $\tilde{c}_3$ in $\mathcal{R}_1$ and $\mathcal{R}_3$. It is often (as it is here) assumed that $A$ and $B$ are available in such a large amount that they are practically constant. Consequently, the associated stoichiometry matrix is

$$\mathbf{S} = \begin{bmatrix}1 & -1 & 1 & -1\end{bmatrix}^T \tag{S26}$$

and the propensity (hazard) vector is

$$\mathbf{h}(X_t, \mathbf{c}) = \begin{bmatrix}h_1, & h_2, & h_3, & h_4\end{bmatrix}^T = \begin{bmatrix}c_1 X(X-1), & c_2 X(X-1)(X-2), & c_3, & c_4 X\end{bmatrix}^T. \tag{S27}$$

Here, since $A$ and $B$ are assumed constant, we have that $c_1 = \tilde{c}_1 A$ and $c_3 = \tilde{c}_3 B$. For the Langevian-approximation, the drift-vector and diffusion matrix are given by

$$\alpha = \begin{bmatrix}h_1 - h_2 + h_3 - h_4\end{bmatrix}, \quad \beta = \begin{bmatrix}h_1 + h_2 + h_3 + h_4\end{bmatrix}. \tag{S28}$$

The Schlögl model is a toy-model, however it exhibits stochastic bi-stability. Hence, it is a suitable benchmark example to use in order to investigate if our framework can infer parameters for challenging stochastic biological systems.

##### S5.3.1 Simulation details

Using Gillespie's direct method, 40 individuals were simulated from the Schlögl model. For each individual, we simulated data at 99 equidistant time-points from 1 to 50 minutes. Regarding observations, we assumed that $X_t$ was directly observed and noise-corrupted by a Gaussian additive error, $Y_t = X_t + \epsilon_t$ where $\epsilon_t \sim \mathcal{N}(0, 2^2)$. The parameters $\mathbf{c} = (c_1, c_2, c_3, c_4)$ were divided into parameters that vary between individuals, here $c_1$, and those constant between individuals, here $\boldsymbol{\kappa} = (c_2, c_3)$. When simulating data, $c_4$ was set at $c_4 = 37.5$ and $\boldsymbol{\kappa} = (-2.17, -8.73)$. The varying parameter $c_1^{(i)}$ were simulated from a log-normal distribution as $c_1^{(i)} \sim \mathcal{LN}(\mu, \tau^2)$ with $\mu = 7.2$ and $\tau = 0.1$.

##### S5.3.2 Inference details

The parameters inferred where the individual parameters $\phi^{(i)} = c_1^{(i)}$, the population parameters $(\mu, \tau)$, the between individuals constant parameters $\boldsymbol{\kappa}$ and the measurement error standard deviation $\sigma$. A weak $\mathcal{N}(0, 10)$ prior was employed for $\mu$. For $\tau$ we used a half-Cauchy prior $\tau \sim \mathcal{C}(0, 2.5)$ where $\tau > 0$. As $\boldsymbol{\kappa}$ was inferred on the log-scale, we used $\log(\kappa) \sim \mathcal{N}(0, 10)$. Lastly, we used a gamma$(2, 1)$ prior for $\sigma$.

We ran the inference using the modifed Gibbs-sampler ($\boldsymbol{\kappa}, \boldsymbol{\xi}$ cell-varying). As the mean of the priors yielded an non-finite likelihood we used a random sample from the prior as starting value. The inference was run according to the guidelines in S3. Briefly, for the pilot run 10 000 iterations were performed using 1000 particles. The high number of particles were employed to control the variability of the likelihood. To further control the variability of the likelihood we used a guided particle filter (modified diffusion bridge), and we correlated the particles with a correlation level $\rho = 0.999$. At the end location of the pilot run, according to the criteria for correlated particle filters, we tuned the number of particles, and the minimum, median and maximum particles required for an individual was 10, 10 and 20 respectively. Lastly, starting from where the pilot-run ended, 500 000 further iterations of the sampler were produced.

#### S5.4 Ornstein-Uhlenbeck model

The Ornstein–Uhlenbeck (OU) is a stochastic differential equation (SDE) model

$$\mathrm{d}X_t^{(i)} = c_1^{(i)}(c_2^{(i)} - X_t^{(i)})\mathrm{d}t + c_3^{(i)}\mathrm{d}W_t^{(i)}, \tag{S29}$$

where the $dW_t^{(i)}$ are increments of standard Brownian motions.

The Ornstein-Uhlenbeck (OU) model is a suitable benchmark model for state-space mixed-effects (SSMEM) framework which, for the case where SSMEMs have latent dynamics $\{X_t^{(i)}\}$ driven by SDEs, it has been named SDEMEMs . A list of resources for inference for SDEMEMs is available at `https://umbertopicchini.github.io/sdemem/`. The process solution to Eq. S29 has a known Gaussian transition density. The fact that Eq. S29 is linear in the latent states implies that if we assume that the observed data follows a linear error model with additive Gaussian measurement noise, $Y_t^{(i)} = X_t^{(i)} + \epsilon_t^{(i)}$, where $\epsilon_t \sim \mathcal{N}(0, \sigma^2)$, then the observed data likelihood (Eq. S3) can be evaluated exactly using a Kalman filter [S41]. Consequently, exact Bayesian inference for the OU model can be performed by plugging the Kalman-provided observed likelihood into our Gibbs sampler. As the Kalman approach does not rely on stochastic approximations of the likelihood, it is a gold-standard benchmark example. Here, we use observed data, and inference results from [S2] when embedding the Kalman-filter in the Gibbs-sampler. In [S2] they produced inference by using the stochastic differential mixed-effects framework that our framework is based on.

##### S5.4.1 Simulation details

The OU-data was generated following [S2]. Briefly, data was simulated for 40 individuals at 200 equidistant time-points from $t = 0.05$ to $t = 10.0$. Regarding the observations, it was assumed that $X_t$ was corrupted by Gaussian additive noise; $Y_t = X_t + \epsilon_t$ where $\epsilon_t \sim \mathcal{N}(0, 0.3^2)$. The individual parameters were generated generated on the log-scale. Here, it was assumed that log-parameters $\boldsymbol{\phi}^{(i)} = \log(\mathbf{c}^{(i)})$ followed a normal-distribution; $\boldsymbol{\phi}^{(i)} \sim \mathcal{N}(\boldsymbol{\mu}, \boldsymbol{\tau}^{-1})$, with $\boldsymbol{\mu} = (-0.7, 2.3, -0.9)$ and $\boldsymbol{\tau} = (4, 10, 4)$.

##### S5.4.2 Validation of the inference framework

To test the correctness of the implementation of our framework, we ran PEPSDI with the non-perturbed option ($\boldsymbol{\kappa}, \boldsymbol{\xi}$ are constant between cells) and the default perturbed option ($\boldsymbol{\kappa}, \boldsymbol{\xi}$ vary weakly between cells) for the OU-model and compared with the exact inference provided by using the Kalman filter to compute the likelihood function and embedding the latter in a Gibbs sampler. PEPSDI performs well (Fig S5). There is some bias for $\tau_2$, however for all other parameters PEPSDI performs well.

Overall, PEPSDI with the default perturbed option correctly infers the population parameters $(\boldsymbol{\mu}, \boldsymbol{\tau})$. The only drawback is a slightly wider credibility interval for that one parameters that is constant between individuals ($\sigma$), however differences in this case are minimal, to the second decimal digit.

### S6 Model of Mig1 nuclear dynamics

The Mig1-model aims to describe the dynamics of the ratio between nuclear Mig1 and cytosolic Mig1 (Mig1n/Mig1c) upon fructose addition to carbon starved cells. To calibrate the model we used two single-cell time-lapse datasets (Fig. 5) where the external fructose concentration was elevated from 0% to 0.05% and 2%, respectively, at t=1.5 min after the start of images acquisition. This results in a rapid increase in the ratio, followed by a decrease corresponding to the Mig1 nuclear entry followed by relocation to the cytosol. The extend of the cytosolic fraction of Mig1 depends on the fructose concentration.

To deduce the mechanisms governing the Mig1-dynamics we considered two pathway structures (Fig. 6a-b). A biological explanation behind each model is provided in the main text. Since model-structure 1 (Fig. 6a) is a sub-structure of model-structure 2 (Fig. 6b) we here focus on model-structure 2.

Overall, model-structure 2 consists of eight reactions:

$$\begin{aligned}
\mathcal{R}_1 &: Frc^{ext} \rightarrow Reg1 \\
\mathcal{R}_2 &: Frc^{ext} \rightarrow M \\
\mathcal{R}_3 &: Mig1c + Reg1 \rightarrow Mig1n + Reg1 \\
\mathcal{R}_4 &: M \rightarrow \phi \\
\mathcal{R}_5 &: M + Mig1n \rightarrow Mig1c + M \\
\mathcal{R}_6 &: Mig1n \rightarrow Mig1c \\
\mathcal{R}_7 &: Mig1c \rightarrow Mig1n \\
\mathcal{R}_8 &: Reg1 \rightarrow \phi
\end{aligned} \tag{S30}$$

Here $M$ refers to the metabolic component. Noticeably $Reg1$ and $M$ are not consumed in $(\mathcal{R}_3, \mathcal{R}_5)$. The is because Reg1 is a phosphatase, which likely acts on Mig1 by dephosphorylation [S42]. Further, $M$ thus likely has phosphate/kinase activity.

Overall, there are eight reactions with associated rate-constants $(c_1, \ldots, c_8)$. Below, each reaction is outlined in detail, and the choice of the prior for each parameter is motivated. The reactions characteristics are further outlined in Tab. S1. Following this, the initial values used for each state are described.

#### S6.1 Motivation of model reactions

$\mathcal{R}_1$ describes the activation of Reg1 via an external carbon-source. Albeit we know that Reg1 is regulated by carbon source availability [S43], intermediate steps are likely encapsulated by $c_1$. These include the activity of hexose-transporters, and the catalytic ability of hexokinases. As we figured that these factors should vary between cells, we allowed $c_1$ to vary between cells. Moreover, $c_1$ should also depend on the external fructose level. The most simple explanation is a proportional dependence. For a typical cell $c_1^{cell} = c_1 * [Fructose]$, however such a proportional scaling fails to properly describe the observed data. Thus, we allowed the log-mean value of $c_1$, $\mu_1$, to vary dependent on fructose availability. For $[Fructose] = 0.05\%$ the log-mean value of $c_1$ is thus different to when $[Fructose] = 2.0\%$.

$\mathcal{R}_8$ describes the degradation of Reg1. For the simplicity of modelling, we neglect the multiple intermediate breakdown steps of Reg1 that we do not think contribute substantially to the cell-to-cell variability and therefore assume that $c_8$ is constant between cells.

For $c_1$ and $c_8$ we use informative priors. Typically, the amount of Reg1 per cell is of the magnitude of several thousands [S44]. Moreover, when adding glucose to starved cells it takes around 3.5 minutes for the Mig1-ration to peak and plateau [S45, S46], and we expect a similar response time when adding fructose. As we expect $M$ to be a delayed signal, due to the observed long-term nuclear export, it thus follows that Reg1 should in 2% fructose reach a peak in around 3.5-minutes. If we now assume that peak amount Reg1 per cell is around 2000 (the amount Reg1 per cell is of $\mathcal{O}(10^3)$) it thus follows that $c_1 \approx 1500$, and $c_8 \approx 0.75$ (see below). Here, $c_1$ applies for [fru] = 2.0%. Since we let $c_1$ vary between cells we employ the prior $\log \mu_1^{(Fru=2.0)} \sim \mathcal{N}(7.3, 2^2)$ and $\log \mu_1^{(Fru=0.05)} \sim \mathcal{N}(6.9, 2^2)$, where $\mathcal{N}$ refers to the normal-distribution. The latter prior highlights that we expect a weaker response, that is activation of Reg1, for lower fructose concentrations [S45, S47].

$\mathcal{R}_6, \mathcal{R}_7$ describes the carbon source independent shuttling of Mig1 [S45]. As the cells at time zero, when no carbon source in the environment is available, are at a steady state [S45], we have that $c_7 = c_6 Mig1n(t_0)/Mig1c(t_0)$. With the same argument as for $c_8$ we assume that $c_6$ is cell-constant ($c_7$ is almost constant as $Mig1n_{t_0} \ll Mig1c_{t_0}$). Moreover, since the Mig1 shuttling, on our timescale, is fast with a FRAP recovery time around 5 seconds [S45], we fixed $c_6 = 40.0$ (see below). We know that the "true" value is likely not 40, however this value encapsulates, on our time-scale, fast nuclear shuttling.

$\mathcal{R}_2, \mathcal{R}_4$ describe the activation and degradation of $M$. As with $c_1$, we expect that multiple processes are encapsulated by $c_2$. Consequently, we let $c_2$ vary between cells. Moreover, similarly to $c_1$ we assume that the log-mean value of $c_2$, denoted $\mu_2$, differs depending on fructose availability. As with $c_8$, we assume that $c_4$ is constant between cells. Our prior knowledge regarding $(c_2, c_4)$ is limited. However, we know that the growth rate of M should be relatively slow (as the feedback is slow, due to the delayed signal (see above for Reg1)). Consequently, we put a prior $\log \mu_2 \sim \mathcal{N}(2, 3^2)$ for both $[Fru] = 2.0$ and $[Fru] = 0.05$. For the scale-parameter of $c_2$ we employed half-Cachy priors $\tau \sim \mathcal{C}(0, 2.5)$ where $\tau > 0$. Further, we put $\log c_4 \sim \mathcal{N}(-6, 3^2)$. The latter ensures that M does not plateau, and the amount of M is relatively small (around 100), thus we are able to investigate if the intrinsic noise might play a large role in the cellular dynamics. To compensate for a wrong assumptions regarding $M$, we employ relatively weak priors.

$\mathcal{R}_5$ describes the $M$ dependent shuttling of into the cytosol. As we assume M is around 100 we know that $c_3$ should be in a range of about 0.05 to 20 (else all Mig1 will leave the nucleus), thus we used a $\log c_5 \sim \mathcal{N}(0, 3^2)$ prior.

Overall, we have that $c_1, c_2$ are cell-varying. The remaining rate-constants $c_{3:8} = (c_3, \ldots, c_8)$ likely vary between cells. However, since we also have variability in the model components (e.g varying Reg1 amount between cells) and $c_{3:8}$ describes fewer cellular processes compared to $(c_1, c_2)$, that is should vary less between cells, we here assumed negligible extrinsic variability.

#### S6.2 Initial values

The model consists of four states: $(Reg1, Mig1c, Mig1n, M)$. As we assume that $Reg1$ and $M$ are activated by the external carbon-source, their initial values are set to zero. Naturally, the number of active Reg1 molecules are not zero, however, since Reg1 does not interact with Glc7 in low carbon source conditions, we expect the amount of active Reg1 molecules to be low [S48].

The initial amount of Mig1c and Mig1n molecules are not zero. Based on available numbers [S47], we know that $Mig1c_{t_0}$ is around 1500 and $Mig1n_{t_0}$ is around 100 molecules per cell when

**Table S1:** Reactions included in Mig1-model

| | Description | Propensity | Comment |
|---|---|---|---|
| $\mathcal{R}_1$ | Activation of Reg1 via Fructose | $c_1$ | $c_1$ cell varying |
| $\mathcal{R}_2$ | Activation of M via Fructose | $c_2$ | $c_2$ cell varying |
| $\mathcal{R}_3$ | Reg1 dependent transport of Mig1 into nucleus | $c_3 Mig1c * Reg1$ | $c_3$ cells constant |
| $\mathcal{R}_4$ | Degradation of M | $c_4 M$ | $c_4$ cells constant |
| $\mathcal{R}_5$ | M dependent transport of Mig1 into cytosol | $c_5 Mig1n * M$ | $c_5$ cells constant |
| $\mathcal{R}_6$ | Frc independent transport of Mig1 into cytosol | $c_6 Mig1n$ | $c_6 = 40$ |
| $\mathcal{R}_7$ | Frc independent transport of Mig1 into nucleus | $c_7 Mig1c$ | $c_7 = c_6 \frac{Mig1n_{t_0}}{Mig1c_{t_0}}$ |
| $\mathcal{R}_8$ | Degradation of Reg1 | $c_8 Reg1$ | $c_8$ cells constant |

glucose or fructose are not available. Moreover, the Mig1 amount should vary between cells [S45, S46]. Consequently, for the initial values we employed priors $\log Mig1c_{t_0} \sim \mathcal{N}(7.3, 0.2)$ and $\log Mig1c_{t_0} \sim \mathcal{N}(4.6, 0.2)$. Furthermore, we know from previous experiments that the variance of $\log Mig1c_{t_0}$ is around 0.2 [S1]. Hence, on the scale component of $Mig1_{t_0}$ we employed $\mathcal{G}(0.4, 0.4)$ priors.

Overall $(c_1, c_2, Mig1n_{t_0}, Mig1c_{t_0})$ vary between cells. As we suspected a relatively strong correlation (e.g between Mig1 in the cytosol and nucleus) we employed a LKJ(0.1) prior on the correlation matrix $\mathbf{\Phi}$.

#### S6.3 Derivation of priors and quantity values

To fix the value of $c_6$ to 40 we simulated a fluorescence recovery after photobleaching (FRAP) experiment. More specifically, to emulate the setting in [S45] we consider a scenario where Mig1 is GFP-tagged, no external carbon source is available, and the nucleus is bleached. Specifically, we consider the nuclear shuttling reactions

$$\begin{aligned} c_6 &: \; Mig1c \rightarrow Mig1n \\ c_7 &: \; Mig1n \rightarrow Mig1c \end{aligned} \tag{S31}$$

with initial values $Mig1c_{t_0} = 1500$ and $Mig1n_{t_0} = 100$ and $c_7 = c_6 Mig1n_{t_0}/Mig1c_{t_0}$. The latter captures that the cells are in a steady state [S45]. To model the FRAP experiment each molecule was given a GFP-tag that equals 1 if the molecule has an active fluorophore, and 0 if it has a none-active fluorophore (i.e. is bleached). At time zero the tag for all molecules in the nucleus was set to zero (to mimic the bleaching), and the system in Eq. S31 was simulated with different values of $c_6$, namely $c_6 = 0.4, 1.4, 2.4, \ldots, 200$. To account for stochastic dynamics each combination was simulated 50 times. By regarding the ratio $Mig1n(GFP = 1)/Mig1n_{tot}$ we noticed that the simulated FRAP-recovery $(Mig1n(GFP = 1)/Mig1n_{tot} \approx 1)$ times decreases with increased $c_6$. Observed recovery time is around 5 seconds [S45] (when the FRAP curve flattens out), and at 1 seconds the recovery is around 0.8. When simulating the reactions in Eq. S31 with different $c_6$-values, this corresponds to a value of around $c_6 \approx 40$.

To obtain priors for $c_1$ and $c_8$ we considered the reaction network

$$\begin{aligned} c_1 &: \; Fru^{ext} \rightarrow Reg1 \\ c_8 &: \; Reg1 \rightarrow \phi \end{aligned} \tag{S32}$$

Our premise here is that Reg1 should start to plateau around $t = 3.5$ minutes, and that the steady state-value should be around 2000 in high fructose conditions ([fru] = 2%). Given this it follows that the steady state value in high fructose should be $c_2 = c_1/2000$. By simulating different different values for $c_1$ using the reactions in Eq. S32, we found that a value of $c_1 = 1500$ fulfils these criteria.

#### S6.4 Inference details

The parameters inferred were the individual parameters $(c_1^{(i)}, c_2^{(i)}, Mig1n_{t_0}^{(i)}, Mig1c_{t_0}^{(i)})$, the cell-constant parameters $c_3, c_4, c_5, c_8$, the strength of the measurement error $\xi$ and the population parameters $\boldsymbol{\eta} = (\boldsymbol{\mu}, \boldsymbol{\tau}, \boldsymbol{\Phi})$. The priors used are outlined above.

Since we did not expect the molecule number in the states $(Reg1, Mig1c, Mig1n, M)$ to be low (e.g 2-10 molecules), we simulated the model dynamics using the tau-leaping simulator with a fixed step length. Due to stiffness, we had to employ a small step length of $\Delta t = 5 \times 10^{-3}$. To achieve efficient inference we correlated the particles with a correlation $\rho = 0.999$. Since we strongly correlated the particles, we could use relatively few particles to estimate the likelihood (see below).

We ran the inference using the default options in PEPSDI, where the cell-constant parameters are slightly perturbed. We used the mean of the priors as starting values. The inference was run according to the guidelines in S3. Briefly, for each model (Model 1A/B and Model 2A/B Fig. 6) we ran multiple pilot runs using 200-300 particles and 1500-3000 samples. At the end location of the pilot run, we tuned the number of particles per individual according to the criteria for non-correlated particle filters. Lastly, starting from where the pilot-run ended, 40,000 further iterations of the perturbed Gibbs sampler (default option in PEPSDI) were run to produce the final inference.

### References Supplementary

[S1] Schmidt GW, Welkenhuysen N, Ye T, Cvijovic M, Hohmann S. Mig1 localization exhibits biphasic behavior which is controlled by both metabolic and regulatory roles of the sugar kinases. Molecular Genetics and Genomics. 2020 nov;295(6):1489–1500.

[S2] Wiqvist S, Golightly A, McLean AT, Picchini U. Efficient inference for stochastic differential equation mixed-effects models using correlated particle pseudo-marginal algorithms. Computational Statistics and Data Analysis. 2021 may;157:107151.

[S3] Hoffman MD, Gelman A. The No-U-Turn Sampler: Adaptively Setting Path Lengths in Hamiltonian Monte Carlo; 2014.

[S4] Ge H, Xu K, Ghahramani Z. Turing: a language for flexible probabilistic inference. In: International Conference on Artificial Intelligence and Statistics, {AISTATS} 2018, 9-11 April 2018, Playa Blanca, Lanzarote, Canary Islands, Spain; 2018. p. 1682–1690.

[S5] Xu K, Ge H, Tebbutt W, Tarek M, Trapp M, Ghahramani Z. AdvancedHMC.jl: A robust, modular and efficient implementation of advanced HMC algorithms; 2019.

[S6] Gillespie DT, Hellander A, Petzold LR. Perspective: Stochastic algorithms for chemical kinetics. Journal of Chemical Physics. 2013;138(17).

[S7] Gillespie DT. Exact stochastic simulation of coupled chemical reactions. In: Journal of Physical Chemistry. vol. 81. American Chemical Society; 1977. p. 2340–2361.

[S8] Voliotis M, Thomas P, Grima R, Bowsher CG. Stochastic Simulation of Biomolecular Networks in Dynamic Environments. PLOS Computational Biology. 2016 jun;12(6):e1004923.

[S9] Gillespie DT. Chemical Langevin equation. Journal of Chemical Physics. 2000 jul;113(1):297–306.

[S10] Golightly A, Gillespie CS. Simulation of stochastic kinetic models. Methods in Molecular Biology. 2013;1021:169–187.

[S11] Haseltine EL, Rawlings JB. Approximate simulation of coupled fast and slow reactions for stochastic chemical kinetics. Journal of Chemical Physics. 2002;117(15):6959–6969.

[S12] Cao Y, Gillespie DT, Petzold LR. Efficient step size selection for the tau-leaping simulation method. Journal of Chemical Physics. 2006 jan;124(4):044109.

[S13] Andrieu C, Doucet A, Holenstein R. Particle Markov chain Monte Carlo methods. Journal of the Royal Statistical Society: Series B (Statistical Methodology). 2010 jun;72(3):269–342.

[S14] del Moral P. Feynman-Kac formulae. Genealogical and interactin g particle systems, with applications. Springer Verlag New York, Series {\S}} Probability and its Applications; 2004.

[S15] Pitt MK, Silva RDS, Giordani P, Kohn R. On some properties of Markov chain Monte Carlo simulation methods based on the particle filter. In: Journal of Econometrics. vol. 171. North-Holland; 2012. p. 134–151.

[S16] Naesseth CA, Lindsten F, Schön TB. Elements of Sequential Monte Carlo. Foundations and Trends in Machine Learning. 2019 mar;12(3):187–306.

[S17] Stewart L, McCarty, Jr P. Use of Bayesian belief networks to fuse continuous and discrete information for target recognition, tracking, and situation assessment</title>. Signal Processing, Sensor Fusion, and Target Recognition. 1992 jul;1699(9):177–185.

[S18] Gordon NJ, Salmond DJ, Smith AFM. Novel approach to nonlinear/non-gaussian Bayesian state estimation. IEE Proceedings, Part F: Radar and Signal Processing. 1993;140(2):107–113.

[S19] Durham GB, Gallant AR. Numerical techniques for maximum likelihood estimation of continuous-time diffusion processes. Journal of Business and Economic Statistics. 2002;20(3):297–338.

[S20] Golightly A, Wilkinson DJ. Bayesian inference for stochastic kinetic models using a diffusion approximation. Biometrics. 2005 sep;61(3):781–788.

[S21] Schauer M, Van Der Meulen F, Van Zanten H. Guided Proposals for simulating multi-dimensional diffusion bridges. Bernoulli. 2017 nov;23(4A):2917–2950.

[S22] Voliotis M, Thomas P, Grima R, Bowsher CG. Stochastic Simulation of Biomolecular Networks in Dynamic Environments. PLOS Computational Biology. 2016 jun;12(6):e1004923.

[S23] Whitaker GA, Golightly A, Boys RJ, Sherlock C. Bayesian inference for diffusion-driven mixed-effects models. Bayesian Analysis. 2017 jun;12(2):435–463.

[S24] Botha I, Kohn R, Drovandi C. Particle Methods for Stochastic Differential Equation Mixed Effects Models. Bayesian Analysis. 2020 jun.

[S25] Golightly A, Bradley E, Lowe T, Gillespie CS. Correlated pseudo-marginal schemes for time-discretised stochastic kinetic models. Computational Statistics and Data Analysis. 2019 aug;136:92–107.

[S26] Haario H, Saksman E, Tamminen J. An adaptive Metropolis algorithm. Bernoulli. 2001;7(2):223–242.

[S27] Andrieu C, Thoms J. A tutorial on adaptive MCMC. Statistics and Computing. 2008 dec;18(4):343–373.

[S28] Vihola M. Robust adaptive Metropolis algorithm with coerced acceptance rate. Stat Comput. 2012;22:997–1008.

[S29] Alvarez I, Niemi J, Simpson M. Bayesian inference for a covariance matrix. The Annals of Statistics. 2014 aug;20(4):1669–1696.

[S30] Lixoft. Monolix version 2019R2. Antony, France: Lixoft SAS; http://lixoft.com/products/monolix/. 2019. Available from: `http://lixoft.com/products/monolix/`.

[S31] Davidian M, Giltinan DM. Nonlinear models for repeated measurement data: An overview and update. Journal of Agricultural, Biological, and Environmental Statistics. 2003 dec;8(4):387–419.

[S32] Schmon SM, Deligiannidis G, Doucet A, Pitt MK. Large-sample asymptotics of the pseudo-marginal method. Biometrika. 2021 mar;108(1):37–51.

[S33] Sherlock C, Thiery AH, Roberts GO, Rosenthal JS. On the efficiency of pseudo-marginal random walk metropolis algorithms. Annals of Statistics. 2015 feb;43(1):238–275.

[S34] Doucet A, Pitt MK, Deligiannidis G, Kohn R. Efficient implementation of Markov chain Monte Carlo when using an unbiased likelihood estimator. Biometrika. 2015 jun;102(2):295–313.

[S35] Deligiannidis G, Doucet A, Pitt MK. The Correlated Pseudo-Marginal Method. Journal of the Royal Statistical Society Series B: Statistical Methodology. 2018 nov;80(5):839–870.

[S36] Golightly A, Bradley E, Lowe T, Gillespie CS. Correlated pseudo-marginal schemes for time-discretised stochastic kinetic models. Computational Statistics and Data Analysis. 2019 aug;136:92–107.

[S37] Limpert E, Stahel WA, Abbt M. Log-normal Distributions across the Sciences: Keys and Clues: On the charms of statistics, and how mechanical models resembling gambling machines offer a link to a handy way to characterize log-normal distributions, which can provide deeper insight into v. BioScience. 2001 may;51(5):341–352.

[S38] Limpert E, Stahel WA. Problems with Using the Normal Distribution – and Ways to Improve Quality and Efficiency of Data Analysis. PLoS ONE. 2011 jul;6(7):e21403.

[S39] Carpenter B, Gelman A, Hoffman MD, Lee D, Goodrich B, Betancourt M, et al. Stan : A Probabilistic Programming Language. Journal of Statistical Software. 2017 jan;76(1).

[S40] F Schlögl. Chemical reaction models for non-equilibrium phase transitions. Z Physik. 1972;1972:147–161.

[S41] Särkkä S, Solin A. Applied stochastic differential equations. Cambridge University Press; 2019.

[S42] Shashkova S, Leake MC. Single-molecule fluorescence microscopy review: shedding new light on old problems. Bioscience Reports. 2017 aug;37(4):20170031.

[S43] Castermans D, Somers I, Kriel J, Louwet W, Wera S, Versele M, et al. Glucose-induced posttranslational activation of protein phosphatases PP2A and PP1 in yeast. Cell Research. 2012 jun;22(6):1058–1077.

[S44] Ho B, Baryshnikova A, GW B. Unification of Protein Abundance Datasets Yields a Quantitative Saccharomyces cerevisiae Proteome. Revista de derecho y genoma humano = Law and the human genome review / Catedra de Derecho y Genoma Humano/Fundacion BBV-Diputacion Foral de Bizkaia. 2018 jan;6(2):192–205.e3.

[S45] Bendrioua L, Smedh M, Almquist J, Cvijovic M, Jirstrand M, Goksör M, et al. Yeast AMP-activated protein kinase monitors glucose concentration changes and absolute glucose levels. Journal of Biological Chemistry. 2014 may;289(18):12863–12875.

[S46] Welkenhuysen N, Borgqvist J, Backman M, Bendrioua L, Goksör M, Adiels CB, et al. Single-cell study links metabolism with nutrient signaling and reveals sources of variability. BMC Systems Biology. 2017 jun;11(1):59.

[S47] Wollman AJM, Shashkova S, Hedlund EG, Friemann R, Hohmann S, Leake MC. Transcription factor clusters regulate genes in eukaryotic cells. eLife. 2017 aug;6.

[S48] Rubenstein EM, McCartney RR, Zhang C, Shokat KM, Shirra MK, Arndt KM, et al. Access denied: Snf1 activation loop phosphorylation is controlled by availability of the phosphorylated threonine 210 to the PP1 phosphatase. Journal of Biological Chemistry. 2008 jan;283(1):222–230.
